# “Comparative Analysis of Glycoproteomic Software Using a Tailored Glycan Database”

**DOI:** 10.1101/2024.07.24.604997

**Authors:** Reuben A. Hogan, Lauren E. Pepi, Nicholas M. Riley, Robert J. Chalkley

**Affiliations:** University of California, San Francisco; Beth Israel Deaconess Medical Center, Harvard Medical School; University of Washington

## Abstract

Glycoproteomics is a rapidly developing field, and data analysis has been stimulated by several technological innovations. As a result, there are many software tools from which to choose; and each comes with unique features that can be difficult to compare. This work presents a head-to-head comparison of five modern analytical software: Byonic, Protein Prospector, MSFraggerGlyco, pGlyco3, and GlycoDecipher. To enable a meaningful comparison, parameter variables were minimized. One potential confounding variable is the glycan database that informs glycoproteomic searches. We performed glycomic profiling of the samples and used the output to construct matched glycan databases for each software. Up to 19,000 glycopeptide spectra were identified across three replicates of wild-type SH-SY5Y cells. There was substantial overlap among most software for glycoproteins identified, locations of glycosites, and glycans, although Byonic reported a suspiciously large number of glycoproteins and glycosites of questionable reliability. We show that Protein Prospector identified the most glycopeptide spectrum matches with high agreement to known glycosites in UniProt. Overall, our results indicate that glycoproteomic searches should involve more than one software to generate confidence. It may be useful to consider software with peptide-first approaches and with glycan-first approaches.

## Introduction

Protein glycosylation, the enzymatic attachment of sugars to proteins, is a very heterogeneous yet common post-translational modification (PTM).(1) *N*-glycosylation, which occurs on asparagine residues within a defined motif, is estimated to occur on over half of the proteins encoded in our genome.(2) Not every potential glycosylation site (glycosite) is always occupied. In fact, the specific pattern of occupied glycosites on a given protein, referred to as macroheterogeneity, can indicate its function.(3–5) To make this process even more complex, the sugar chains (glycans) attached at a given glycosite can have different monosaccharide compositions, linkages, and branching structures, referred to collectively as microheterogeneity. This multifaceted heterogeneity makes glycoproteomics, the large-scale study of glycoproteins, challenging.

Modern glycoproteomics is mostly performed using mass spectrometry and requires careful considerations.(6,7) First, one must identify a method to enrich glycopeptides from the background of mostly unmodified peptides.(8) Second, one must select a method of fragmentation that generates informative ions about both peptide and glycan parts of the molecule.(9) Third, one must analyze the spectra in a way that generates confident, reproducible results.(10)

For this final consideration, there are a host of software suites that are available.(1,6) A community evaluation study was published in 2021 to understand how researchers analyzed their data and how consistent their results were.(11) The study showed that manual analysis was able to improve on the best raw search engine outputs, highlighting the room for improvement. Since then, several new software have been developed, and some existing tools have been improved.

A major difference among glycopeptide analysis software is the way in which each software interprets a glycopeptide spectrum. For example, Byonic attempts to identify peptide and glycan components in one step, treating the glycan like a large variable modification.(12,13) It generates a theoretical complete glycopeptide spectrum for all peptide and glycan permutations supplied from user databases and then scores spectra based on their match. Other software, such as Protein Prospector or MSFraggerGlyco, rely on the mass offset approach.(14–16) In this method, masses of potential glycopeptides are calculated in the same way as Byonic, but the fragmentation spectrum is initially only compared to theoretical peptide fragments from the glycopeptide. Having identified the peptide, if there are multiple potential glycans of similar mass, a scoring system is applied to determine the best assignment among these. Newer software, such as pGlyco3 and GlycoDecipher, initially make use of Y-ions.(17,18) Y-ions contain the peptide backbone and fragments of the glycan. Peptide assignment is performed only after the glycan has been identified. Both pGlyco3 and GlycoDecipher flex an ability to identify modified monosaccharides. GlycoDecipher can perform glycan database-independent identification, an effective *de novo* glycan construction. MSFraggerGlyco, pGlyco3, and GlycoDecipher employ false discovery rate (FDR) calculations for both the peptide and the glycan. This could be a great advance in the field over traditional confidence scores, which are difficult to translate into a probability.

Because of the variety of analytical innovations, it is imperative that another benchmark be performed.(19) Publications of the newer software each conducted a comparison to competing tools.(15,17,18,20) However, these only focused on the total number of spectra identified. They did not discuss overlap or agreement and did not break down the results in terms of relevant information such as unique glycopeptides or glycosites. There was also no comparison to the literature to determine agreement with known information.

Herein, we report a comparative benchmarking of five software: Byonic, Protein Prospector, pGlyco3, MSFraggerGlyco, and GlycoDecipher. For this benchmark, we acquired a novel glycoproteomic dataset from SH-SY5Y cells, a human neuroblastoma derived cell line, using strong anion exchange –electrostatic repulsion liquid chromatography (SAX-ERLIC) for glycopeptide enrichment followed by high pH reverse phase fractionation (HpH-RPF). Data was acquired using stepped collisional energy higher energy collisional dissociation (sceHCD), which balances fragmentation quality and acquisition speed to maximize the number of highly quality spectra for *N*-glycopeptide analysis.(9) Of highlight, we performed glycomic profiling of the SH-SY5Y cells to identify the glycans present and constructed matched glycan databases for all searches. Downstream analysis compared search engine results based on multiple criteria, including agreement with reported glycosites.

## Methods

### Cell Culture

SH-SY5Y cells were cultured in 15 cm dishes using DMEM:F12 + 10% FBS with no antibiotics and placed in an incubator at 5% CO_2_ and 98% humidity at 37°C. For subculturing, media was aspirated from the cells. Cells were washed once with PBS before adding accutase. Cells were incubated at 37°C for 5 minutes. Detached cells were collected and centrifuged at 500g for 5 min to pellet in a 1.5 mL microcentrifuge tube. The supernatant was aspirated from the pellet. Pellets were frozen on dry ice and stored at -80°C until lysis. Each replicate used here represents a cell pellet from separate passages.

### Glycomic Profiling

For *N*-glycome profiling, samples were homogenized in an SDS lysis buffer. Proteins were denatured using 10 mM DTT. *N*-glycans were then loaded onto an S-trap plate (Protifi) and incubated with PNGase F at 37°C overnight using the manufacturer’s protocol with minor adjustments for glycomics.(21) Briefly, samples are loaded onto S-trap column using S-trap binding buffer (90% methanol, 100mM TEAB). Samples were eluted from S-trap column with two aliquots of 60 µL 0.1% TFA. Following incubation, *N*-glycans were cleaned by Hypercarb column (ThermoFisher). Hypercarb was conditioned with 2 column volumes (CVs) of 99.9% acetonitrile and 0.1% TFA, followed by 2 CVs of 0.1% TFA. Samples were then loaded onto the columns and washed with 4 CVs of 0.1% TFA. Samples were eluted in 50% acetonitrile with 0.1% TFA. *N*-glycans were analyzed by PGC-LC-MS/MS on a ThermoFisher TSQ Altis Mass Spectrometer coupled to a Vanquish LC system. A targeted *N*-glycan method utilizing over 200 *N*-glycan standards was used for the analysis, and samples were run over an 80 min gradient. Collision energies were previously optimized for each standard. A Dextran ladder was used to normalize retention times across runs.(22) Fragmentation pattern and elution order were compared to the standard library to make glycan assignments. Data was analyzed using ThermoFisher Freestyle software, GlycoWorkbench and Skyline.(23,24)

### Tryptic Digestion

A buffer was made of 8M Urea, 100mM Tris-HCl, 10mM TCEP, 40mM 2-choloroacetamide buffer at pH of 8.0. Frozen cell pellets were lysed in approximately 1mL. Mixture was pipetted up and down until homogenous. Lysate underwent two freeze-thaw cycles. Then, the lysate was sonicated twice by probe tip for 10s at 20% magnitude. Additional cycles were used if the mixture remained viscous. The lysate was incubated at 37°C for 5 min on a Thermomixer at 1500 rpm to reduce cysteines. Lysate was diluted to 1.5M Urea with 100mM Tris-HCl at pH 8.0. Tryspin and Lys-C were added to the lysate at a 1:100 ratio. Digestion ran overnight at 37°C and 1200 rpm.

### Desalting

Following digestion, samples were brought to 1% trifluoroacetic acid (TFA). A Waters Sep-Pac Vac tC18 3cc cartridge was conditioned with three CVs of acetonitrile (ACN) followed by one column volume of 40% ACN / 0.1% TFA. The column was equilibrated with three column volumes of 0.1% TFA. The sample was then loaded onto the column until all the tryptic digest had flown through once. The column was washed with three column volumes of 0.1% TFA. Then, the sample was eluted with 2 mL of 40% ACN / 0.1% TFA followed by 2 mL of 80% ACN / 0.1% TFA. The eluate was lyophilized by SpeedVac.

### SAX-ERLIC Enrichment

This protocol was taken from Bermudez and Pitteri 2021(25). In brief, lyophilized tryptic peptides were resuspended in 1 mL of 50 mM ammonium bicarbonate. The SOLA SAXE SPE cartridge was washed with 3mL of ACN. The column was activated using 3mL of 100mM triethylammonium acetate. The column was conditioned with 3mL of 1% TFA. The column was equilibrated with 3mL of 95% ACN / 1% TFA. Sample was loaded onto the column twice. The column was washed using 6mL of 95% ACN / 1%TFA. Enriched glycopeptides were eluted from the column in two steps: first with two volumes of 850µL of 50% ACN / 1% TFA and second with two volumes of 850uL of 5% ACN / 1% TFA. The two fractions were lyophilized using a SpeedVac.

### High pH-Reverse Phase Fractionation (HpH-RPF)

For HpH-RPF, eight fractions were collected in the following manner. Buffers containing 50%, 20%, 17.5%, 15%, 12.5%, 10%, 7.5%, and 5% ACN in 0.1% triethylamine in water were made. A C18 NEST tip was washed with 200µL of ACN followed by 400µL of 0.1% formic acid. Glycopeptides were resuspended in 100µL of 0.1% formic acid. From 5% ACN to 50% ACN, 100µL of each buffer was added and then eluted by microcentrifuge. All fractions were collected and then lyophilized by SpeedVac. Samples were resuspended in 0.1% formic acid before analysis by mass spectrometry.

### Mass Spectrometry Acquisition

The liquid chromatography gradient was 120 min at constant flow of 600nL/min. Buffer A was 0.1% formic acid. Buffer B was 80% ACN / 0.1% formic acid. Buffer B gradient from 0 to 50% was 113 min followed by a short 5 min 95% Buffer B phase before ending at 0% at 120min.

For Orbitrap Lumos, MS1 resolution was set to 120K. Scan range was 400 to 1800 *m/z*. RF lens was 60%. AGC target was set to 100%. Maximum injection time was 50ms. Dynamic exclusion was set to exclude peaks for 60s after first appearance. Only ion charge states 2-8 were selected. For MS2, fragmentation was performed with sceHCD 20, 30, 40 nce. Resolution was 30K. Scan range was 120 – 2000 *m/z*. AGC target was set to 200%. Maximum injection time was 200ms.

### Data Analysis

All analyses except Protein Prospector were performed on a computer with 1TB RAM and an Intel Xeon 2.40GHz CPU. Protein Prospector was submitted as a job to a server for processing using its web-based interface. All protein databases used the UP000005640_9606 human proteome with one FASTA sequence per protein from UniProt. All glycan databases were informed by glycomic profiling results. For a more detailed explanation, see Glycan Database Conversion. Only specific tryptic peptides with a maximum of 3 missed cleavages were allowed in all searches. All searches allowed for carbamidomethylation as a fixed modification. All searches allowed for oxidation of methionine and protein N-term acetylation as variable modifications.

### Protein Prospector Parameters

Protein Prospector (version 6.5.0) was used for the analysis. Raw data was converted into .mgf format peak list files using in-house software ‘PAVA’, which makes use of Monocle for improved monoisotopic peak selection.(26) These were filtered for the presence of a HexNAc oxonium ion at *m/z* 204.087 (+/- 20ppm) in MSMS scans. The filtered peaklist was searched in Batch-Tag using the same parameters as for other software, other than Gln->pyro-Glu (N-term) was additionally allowed as a variable modification. Also, to adjust for a calibration error in the data, precursor ion mass tolerance considered was a systematic error of 8ppm, then +/- 8 ppm tolerance. Fragment tolerance was 20 ppm. The list of identified glycosylated peptides was then input into MS-Filter to score glycan assignments and find additional glycoforms. Minimum peptide and glycan scores of 0 and 3 were employed, then the best scoring glycan result for each spectrum was reported.

### pGlyco3 Parameters

pGlyco3.1 was run in N-glycan mode. Initial search did not include variable modifications on the glycan and allowed for a glycan database size of 1e5. Subsequent searches allowed for 2 max variable modifications on the glycan with a glycan database size of 1e6. Only peptides between 6 and 40 amino acids long were allowed. Minimum peptide weight was 600Da. Maximum peptide weight was 4000. Carbamidomethylation was allowed as a fixed modification on cysteines. Two max modifications were allowed on the peptide. Precursor tolerance was 10ppm. Glycan and Peptide FDR thresholds were set to 1%.

### MSFraggerGlyco Parameters

FragPipe (v21.1) with “N-glyco-HCD” workflow was loaded as default. Only peptides between 7 and 50 amino acids long were allowed. Three max modifications were allowed on the peptide. Peptide charges between 1 and 4 were considered. Precursor tolerance was 20 ppm. Glycan and Peptide FDR thresholds were set to 1%.

### GlycoDecipher Parameters

GlycoDecipher (v1.0.4) was used. Three max modifications were allowed on the peptide. Peptide length was between 6 and 40 amino acids long. Precursor tolerance was set to 5ppm. Peptide charges between 2 and 6 were considered. Minimum peptide mass was 600Da. Maximum peptide mass was 4500Da.

### Byonic Parameters

Byonic from Protein Metrics (v5.4.52) was used. One max modification was allowed on the peptide. Precursor tolerance was set to 20ppm. Maximum precursor mass was 10000Da.

### MaxQuant Parameters

Peptides were allowed to be between 7 and 40 amino acids long. Maximum peptide mass was 4600Da. Protein FDR was set to 1%.

### Glycan Database Conversion

GlyTouCan Accession Numbers from glycomic profiling were used to import 53 glycan structures into a GlycoWorkbench file. The GlycoWorkbench file was converted into a glycan database file for pGlyco3 (.gdb) using the Convert Glycoworkbench script available in the software. The logic of this script creates all the unique fragments from structures in GlycoWorkbench. This generated a list of 200 unique glycan structures. Custom scripts were created by dictating the logic of conversion to ChatGPT-4 and allowing for it to generate Python scripts that could be run in terminal.(27) Results were manually inspected for accuracy. This approach created glycan databases for Protein Prospector, Byonic, and MSFraggerGlyco. GlycoDecipher uniquely uses the GlyTouCan Accession Numbers. In this case, the contents of the existing “database.csv” file were replaced with only the GlyTouCan Accession Numbers and accessory information.

### Preparation of Results of Glycoproteomic Softwares

Each glycoproteomic software generates results in a different format. Therefore, to perform a comparison, the output required manipulation. For Byonic and GlycoDecipher, ChatGPT-4 was used to create a bash script that would concatenate all the results into a single file.(27) For MSFraggerGlyco, all processing used the “psm.tsv” output. For pGlyco3, the pre-supplied “protein_site_analysis.py” script was used to generate a file that contained all information for downstream processing. For Protein Prospector, a tab-delimited output of the MS-Filter results was used for downstream processing.

### Analysis in R

Once results from all fractions and replicates could be uploaded in single files, analysis was performed completely in R. Logic for processing and specific functions were dictated to ChatGPT-4, which generated scripts that greatly expedited the analysis.(27) For Byonic, results were filtered for presence of a glycan and a peptide score above 200. For MSFraggerGlyco, results were filtered for presence of a single HexNAc as a modification on the peptide. Protein Prospector, pGlyco3 and GlycoDecipher required no additional processing because their outputs only included glycopeptides.

### Cytoscape

UniProt IDs for the glycoproteins identified in all software was searched using the STRING DB plug-in of Cytoscape. Edge information was set to physical protein interactions with a score filter of 0.4. Functional enrichment was performed using the network of proteins against the whole genome.

## Results

### SAX-ERLIC Enrichment With HpH-RPF Produced High Quality Glycopeptide Spectra By All Software

RAW files were searched with MaxQuant to identify non-glycosylated peptides. Although peptide-first search engines identify modified and unmodified peptides, we chose MaxQuant as an independent search engine to determine the quality of the data. It detected 348,623 spectra in total. Then, RAW files were searched using Byonic, Protein Prospector, pGlyco3, MSFraggerGlyco, and GlycoDecipher (**Figure 1A**).The same FASTA protein database and glycan databases were used in each software to minimize software variability. To provide a glycan database specific to the sample, glycomic profiling of SH-SY5Y cells was performed (**Figure 1C**). There were 53 *N*-glycan structures identified (**Supplementary Table 1**). High mannose glycans were the most abundant, which has been a reported feature of neuronal tissue(28).

**Figure 1:**
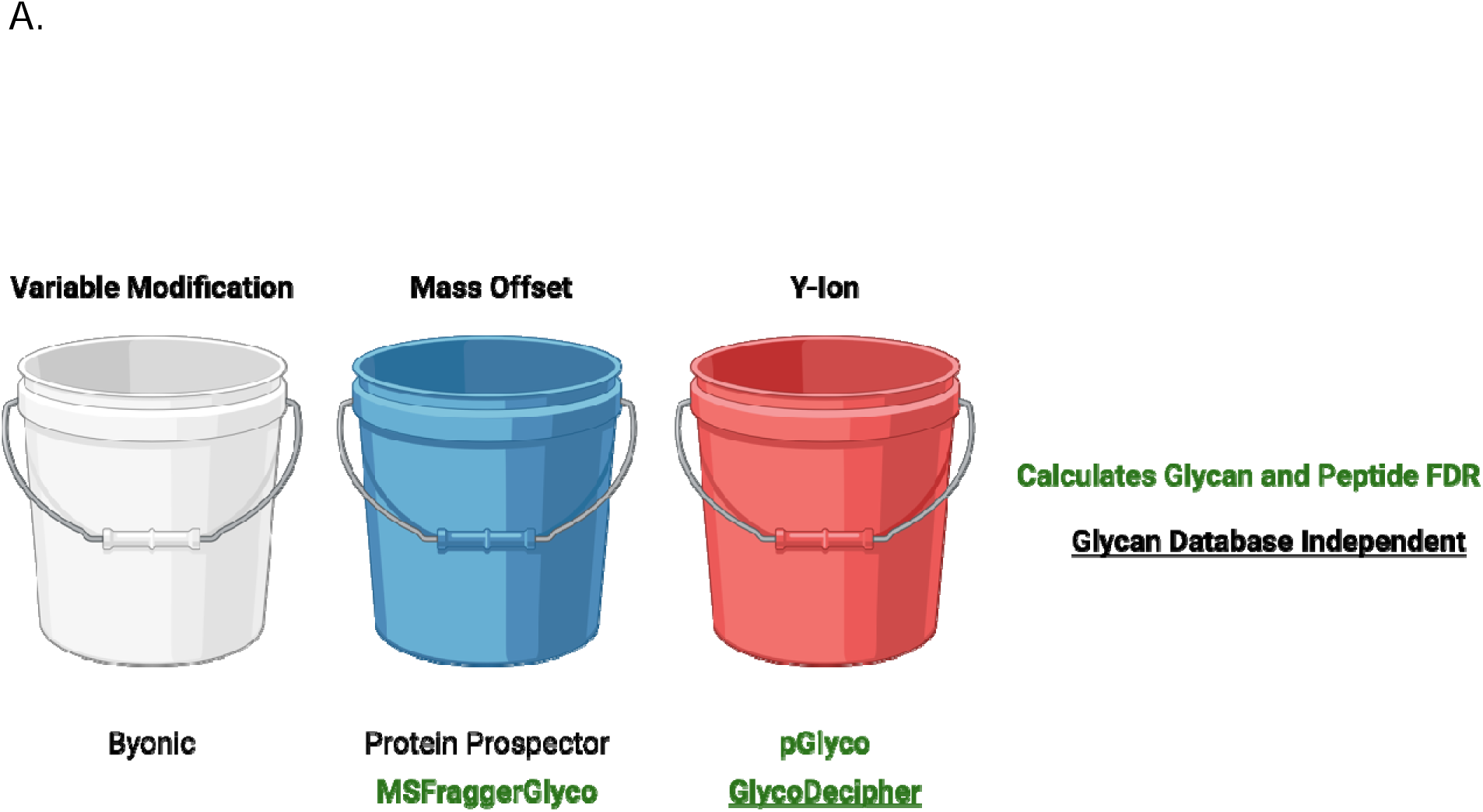
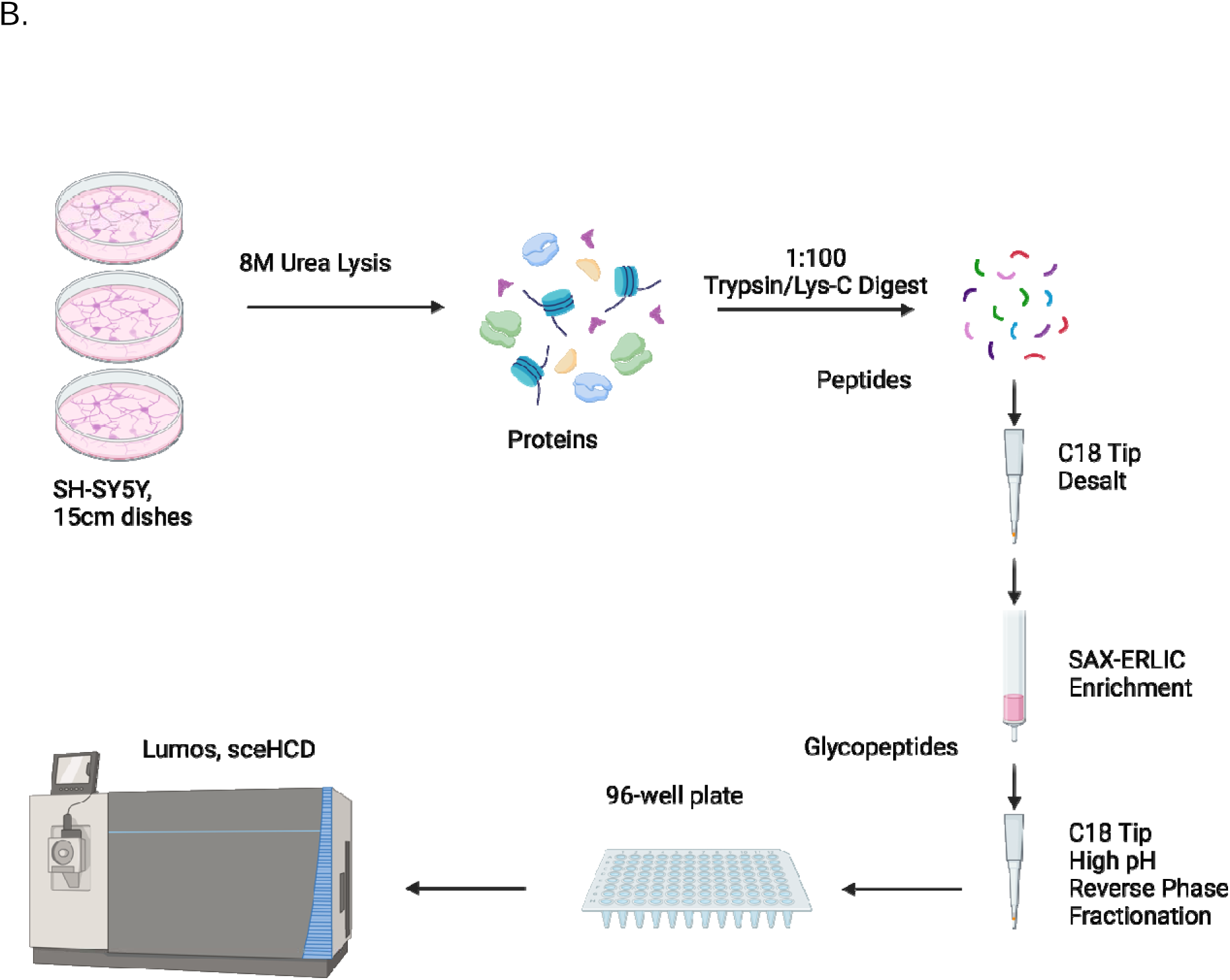
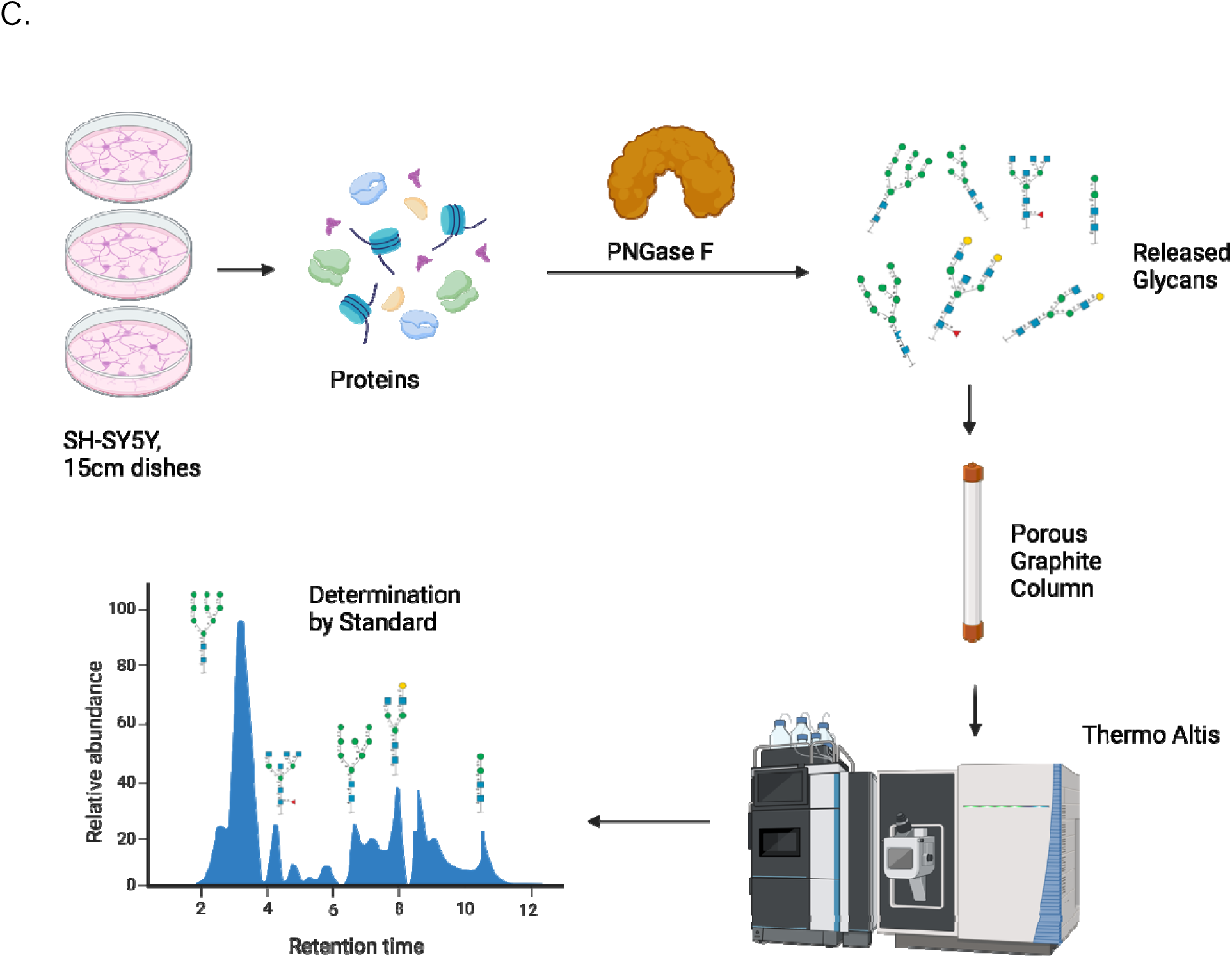
Establishing a Head-to-Head Comparison of Glycoproteomic Softwares. Using Self-Generated Glycoproteomic Data. (A) Layout of the different softwares. (B) Workflow to generate glycopeptides. (C) Workflow for glycomic profiling.

While the number of glycopeptides in a single fraction differed across software, the pattern of glycopeptides found across fractions was consistent (**Supplementary Figure 1**). The most glycopeptides eluted between 10-15% ACN in 0.1% triethylamine. This underscores the hydrophilic nature of most glycopeptides. These plots also indicate that HpH-RPF is effective at separating glycopeptides into fractions of relatively even complexity.

Figure 2A summarizes the number of glycopeptide spectrum matches (GPSMs) reported by each software. In total there were 26,964 unique GPSMs. Protein Prospector reported the most GPSMs (19,717) while pGlyco reported the least (9,482). Other software reported numbers between 11,519 and 14,197. Overall, the mass offset approach software identified the most glycopeptide spectra.

**Figure 2:**
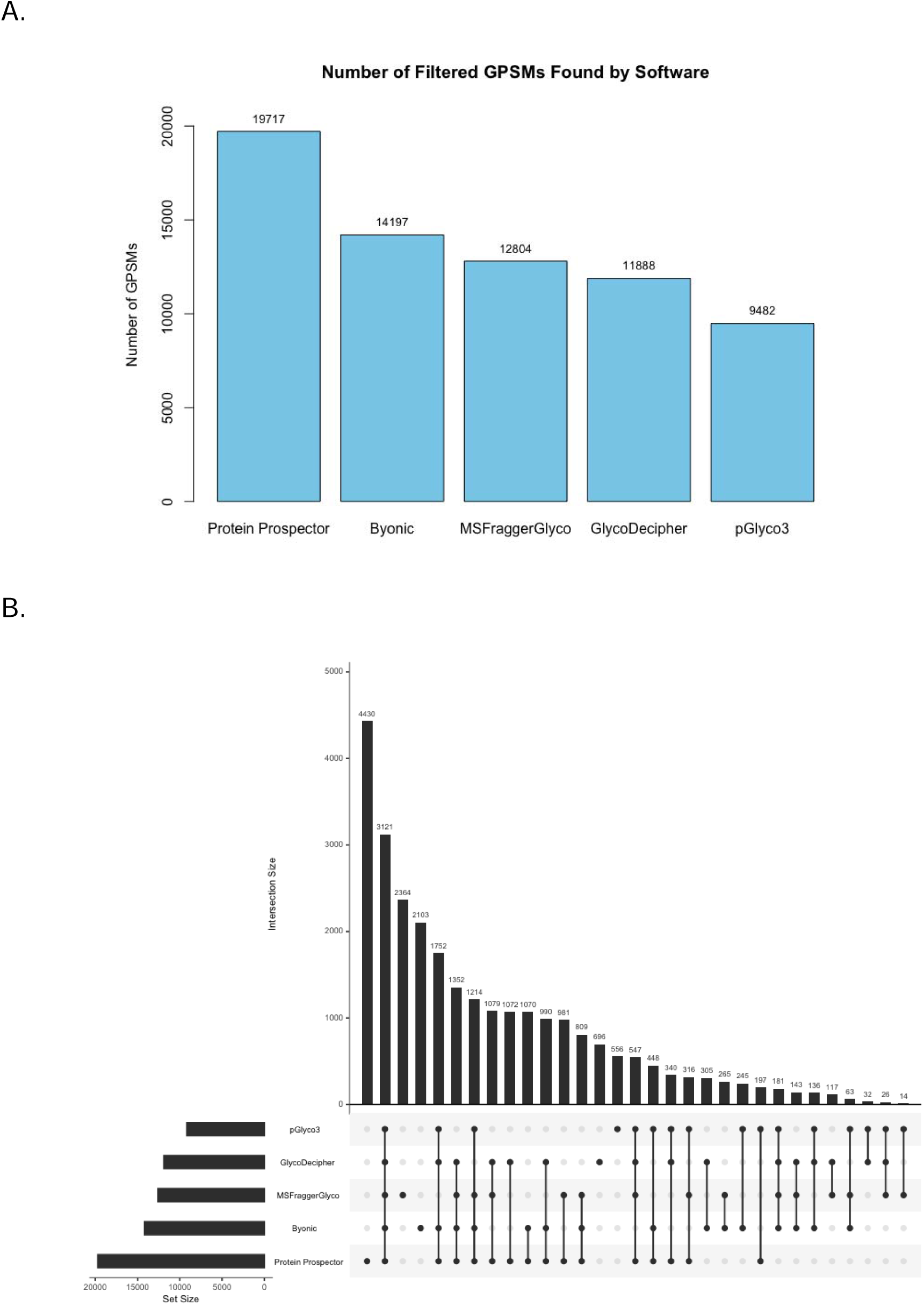
Glycopeptide Spectrum Matches (GPSMs) and Their Overlap. (A) The number of glycopeptide spectral matches that were identified by each software after filtering. (B) UpSet plot of the scan numbers reported for each GPSM.

It is interesting that Protein Prospector identified 5,520 more GPSMs than competing software. To investigate further, we analyzed the scan numbers to determine how many of these were unique spectra and how many were detected by another software (Figure 2B). 4,430 were spectra that Protein Prospector exclusively identified. The remaining 1,090 were spectra that were also assigned in other software. These data suggest that Protein Prospector was more sensitive at identifying glycopeptide spectra. It is possible that this is because Protein Prospector is not using a glycan FDR threshold like MSFraggerGlyco, pGlyco3, and GlycoDecipher.

As an example of what these additional spectra contain, Figure 3A summarizes the results for the peptide AGPNGTLFVADAYK from Adipocyte Plasma Membrane-Associated Protein. Protein Prospector identified more spectra to this peptide than competing software. These covered a total of 10 glycoforms, capturing the second most microheterogeneity among software for this peptide. GlycoDecipher reported more glycoforms from many fewer spectra, although some of these were based on *de novo* assignment of extra glycoforms outside the glycan database. Another example where Protein Prospector identifies more glycoforms than other software is shown in **Supplementary Figure 2**, which shows annotated spectra for additional glycoforms reported for a peptide from Immunoglobulin superfamily member 3.

**Figure 3:**
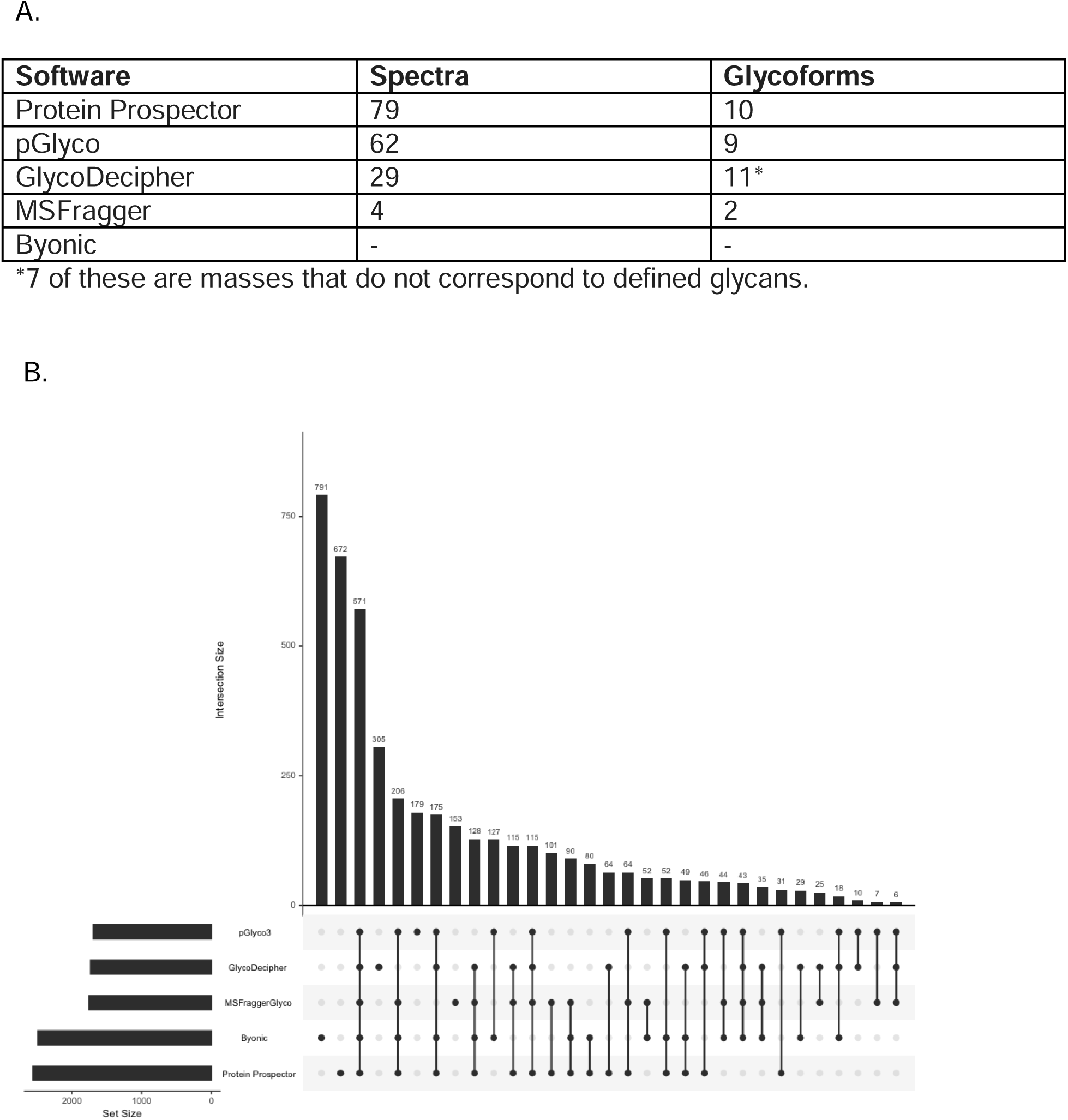
Protein Prospector Captures More Information Content from the Data. (A) Table showing the results from each software for peptide AGPNGTLFVADAYK. (B) UpSet plot of the unique protein-site-glycan combination (PSG IDs) by software.

To understand how many of these spectra corresponded to novel information rather than redundant glycopeptides, we created an identifier for any combination of a protein, site, and glycan (PSG ID). In total, there were 4,383 unique PSG IDs, which represents the total detected diversity of glycopeptides in these samples. Figure 3B shows that Protein Prospector exclusively identified 672 / 4,383 (approximately 15%) PSG IDs. These data indicate that the unique results from Protein Prospector are not redundant information. In fact, Protein Prospector captured the most PSG IDs at 2,559 / 4,383 (approximately 58%) followed by Byonic at 2,490 / 4,383 (approximately 57%).

### Comparison of the Glycoprotein Identities

To understand whether common glycoproteins were identified and how many, we plotted the overlap in results at the protein level (Figure 4A). In total, there were 947 unique glycoproteins found by at least one software. Of these, 231 glycoproteins (approximately 24%) were found by every software, matching the poor overlap seen in the previous community-wide study.(11) A large portion of the unique glycoproteins came from Byonic, which exclusively identified 382 / 947 proteins (approximately 40%). In fact, Byonic was an outlier because it identified a total of 749 glycoproteins where the nearest competitor, MSFraggerGlyco, found a total of 389. By looking at the spectra uniquely identified in Byonic, we were able to determine that many of these were incorrect assignments (for example, see Figure 4B). On the other hand, despite reporting the most GPSMs, Protein Prospector reported the fewest glycoproteins. Excluding Byonic, using at least two software was sufficient to reproduce at least 254 glycoprotein identities. This would be 65% of hits for MSFraggerGlyco or 86% for Protein Prospector.

**Figure 4:**
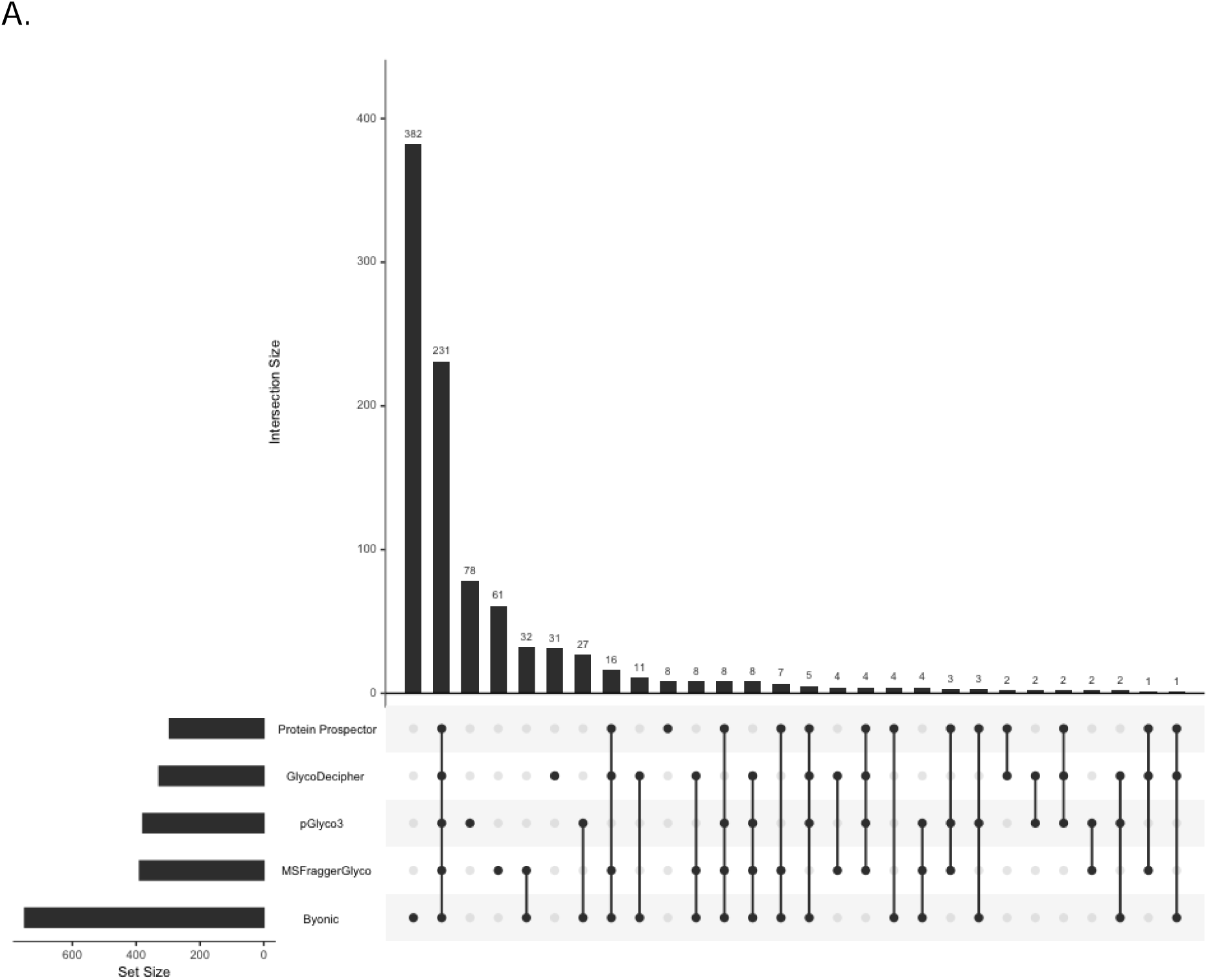
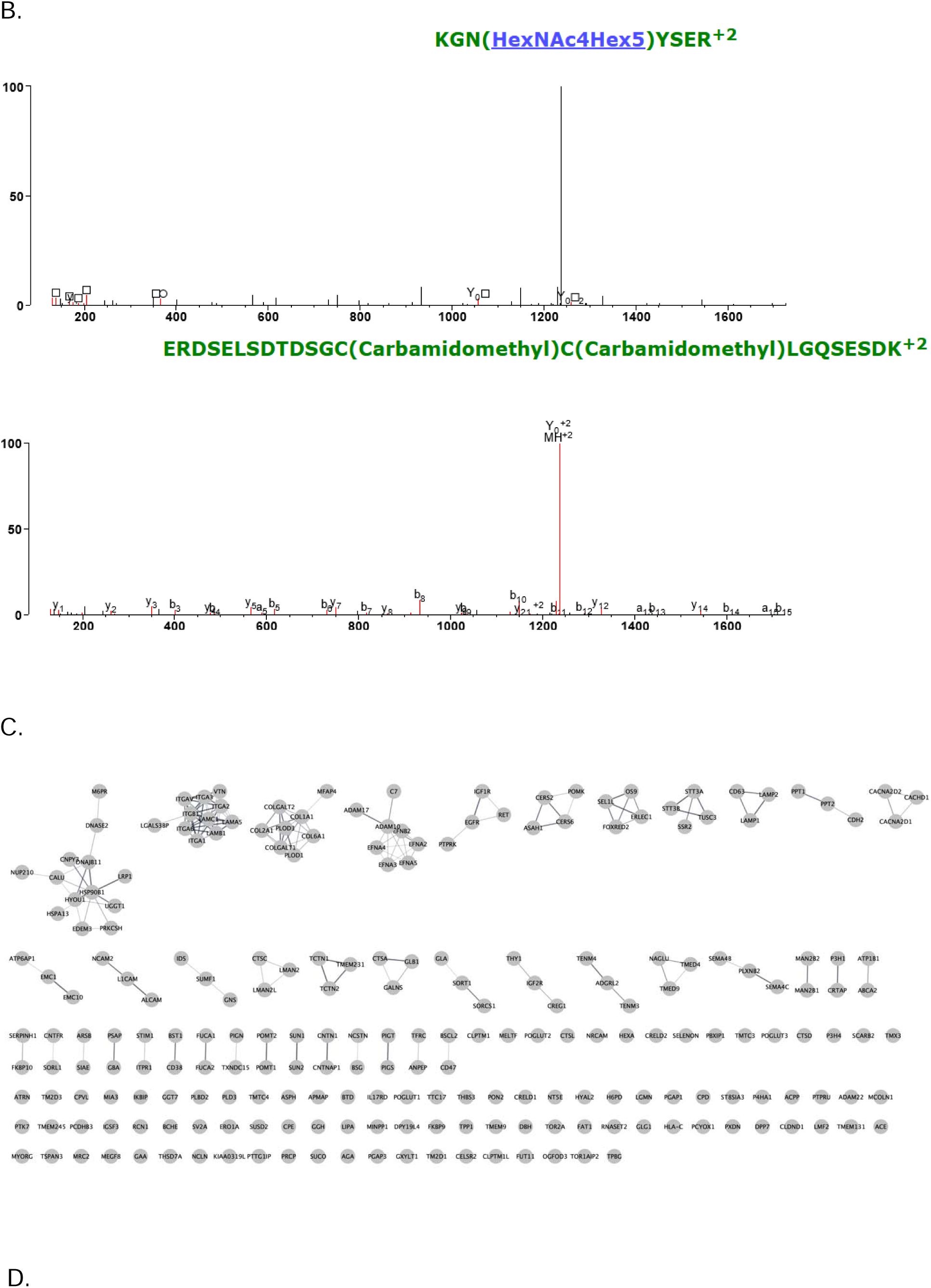
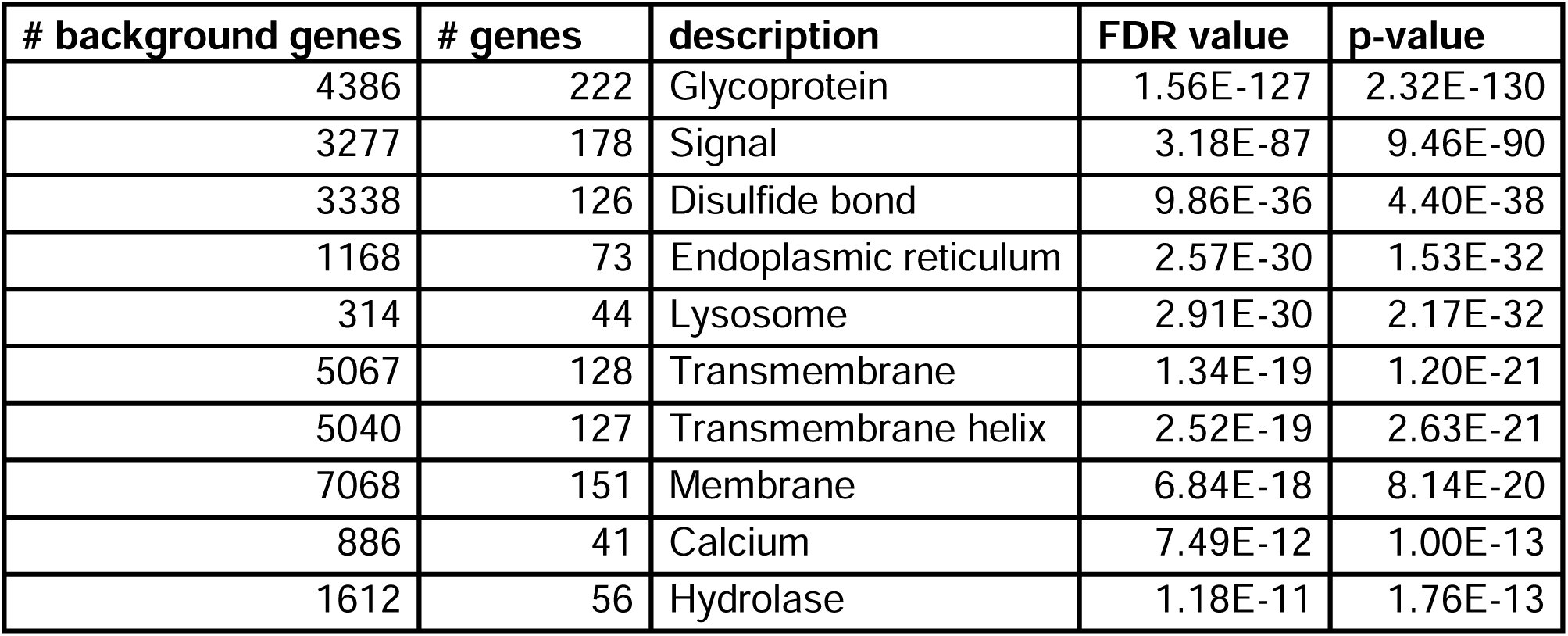
Comparison of the Unique Glycoproteins. (A) UpSet plot of the unique glycoprotein results from each software. (B) This is an example of a unique glycopeptide assignment by Byonic. The peptide Byonic assigns is KGNYSER with glycan HexNAc4Hex5. The peptide comes from HistoneH2A. Protein Prospector assigns this spectrum to an unglycosylated peptide from E3 Ubiquitin-protein ligase UHRF1. (C) Cytoscape network of the common 212 glycoproteins. (D) Functional enrichment for the common glycoproteins.

Encouragingly, there was good confidence in the commonly reported glycoproteins. Using Cytoscape with STRING, a physical interaction network was generated for 231 common glycoproteins. The results were clustered using a granularity level of 4 (Figure 4C). The network was significantly enriched for protein-protein interactions (PPIs) with a reported PPI value of 1.0E-16 with several visible clusters of proteins. Functional enrichment of the network revealed that the top theme from UniProt Keywords categorization was the term “Glycoprotein” (Figure 4D). In comparison, functional enrichment for all peptides identified by MaxQuant did not include “Glycoprotein” at all but rather “Phosphoprotein”, “Acetylation”, and “Cytoplasm” as top themes (**Supplementary Table 1**). These data indicate that for those proteins upon which software agree, there is high confidence in their status as glycoproteins. It is worth noting, however, that the number of results exclusively reported by a single software was considerable (Figure 4A).

### Comparison of Glycosite Assignments

Figure 5A is an UpSet Plot of glycosite assignments reported by software. In total, there were 1,466 unique glycosites discovered in the searches. Of these, 308 (approximately 21%) were common to all software. Again, Byonic was an outlier. It exclusively reported 491 / 1,466 unique glycosites (approximately 33%). 383 of these were because of the glycoproteins it exclusively reported (Figure 4A). Byonic alone reported a total of 1,123 glycosites where the nearest competitor, MSFraggerGlyco, reported 658. As with the glycoprotein-level summary, Protein Prospector reported the fewest glycosites.

**Figure 5:**
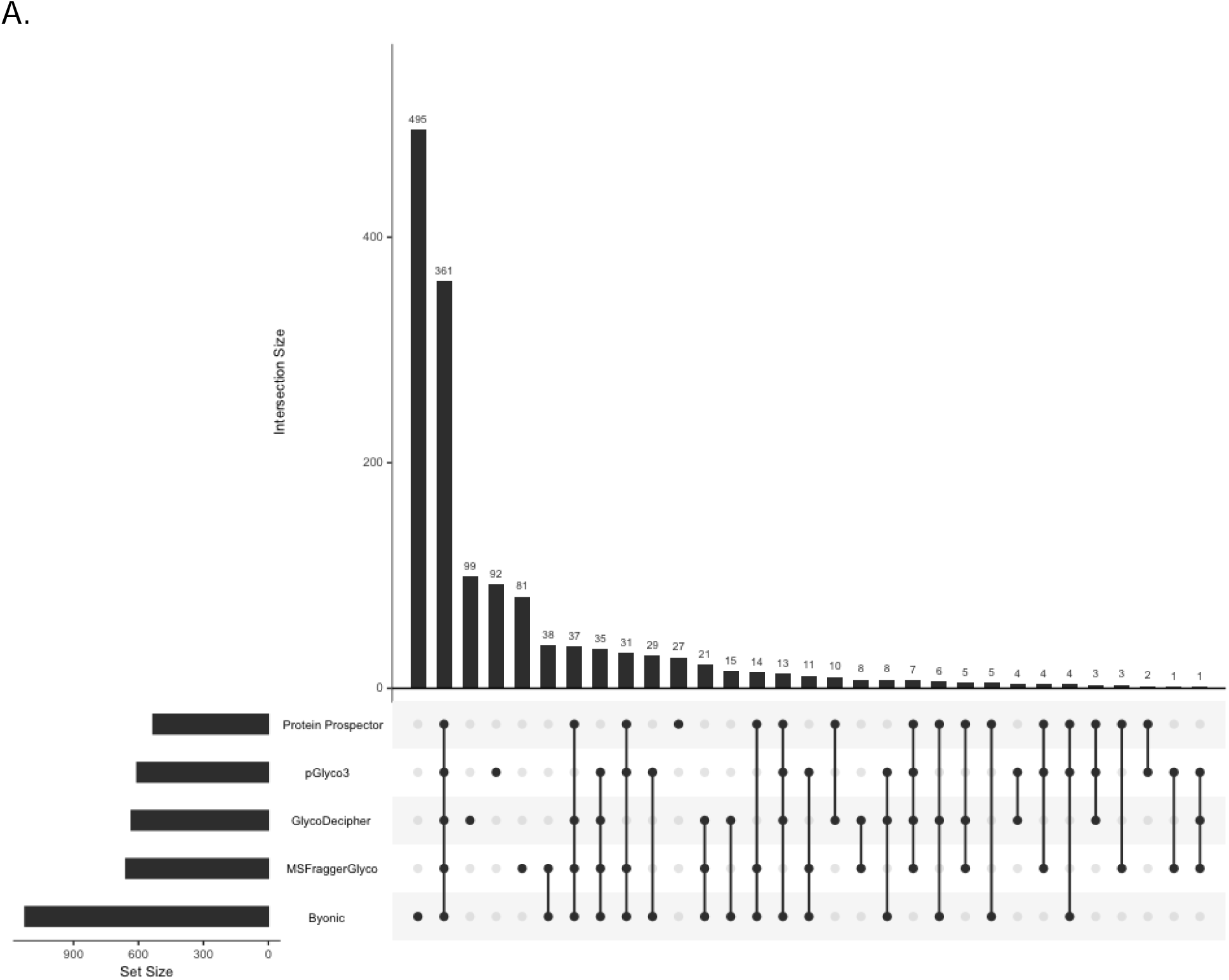
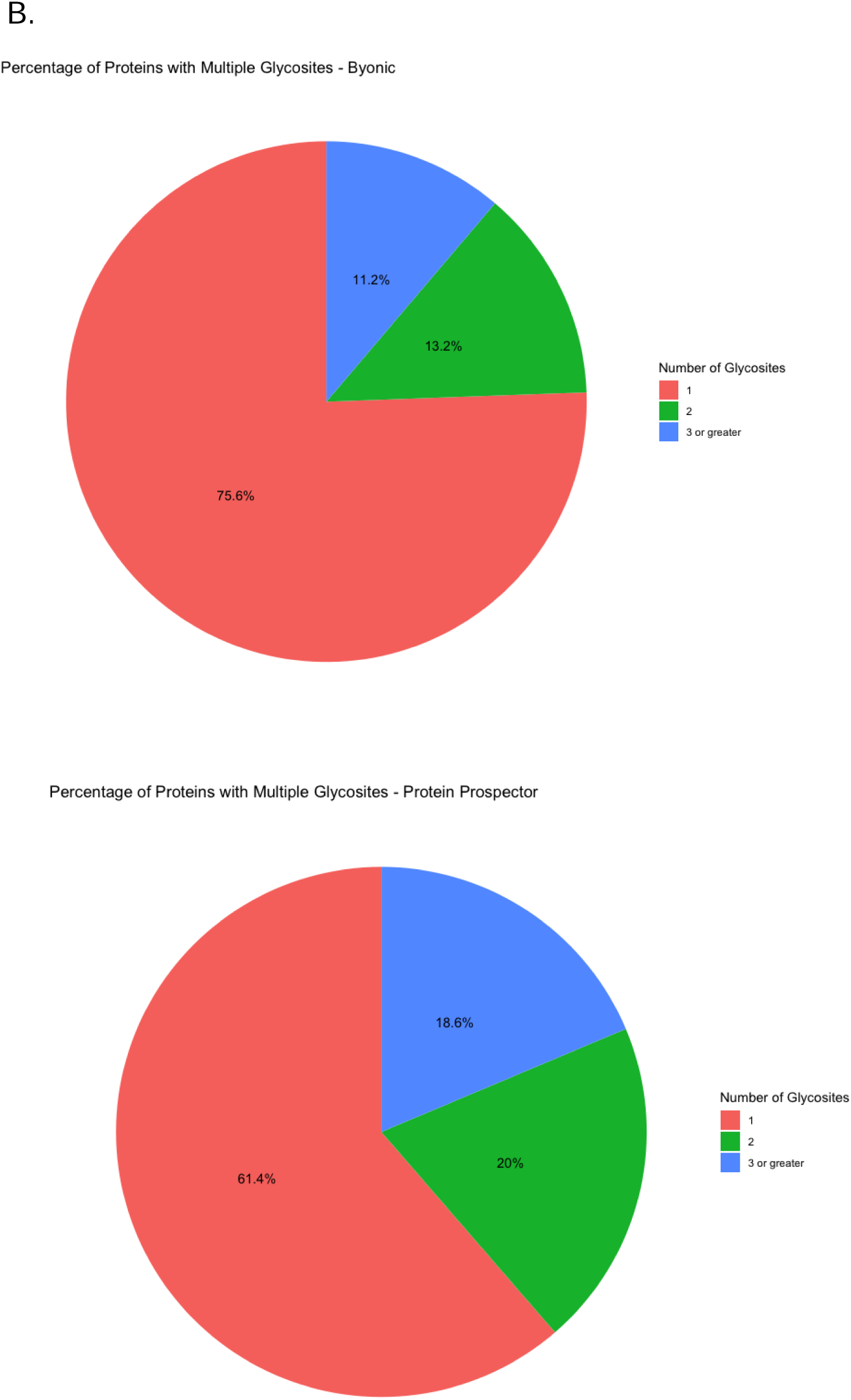
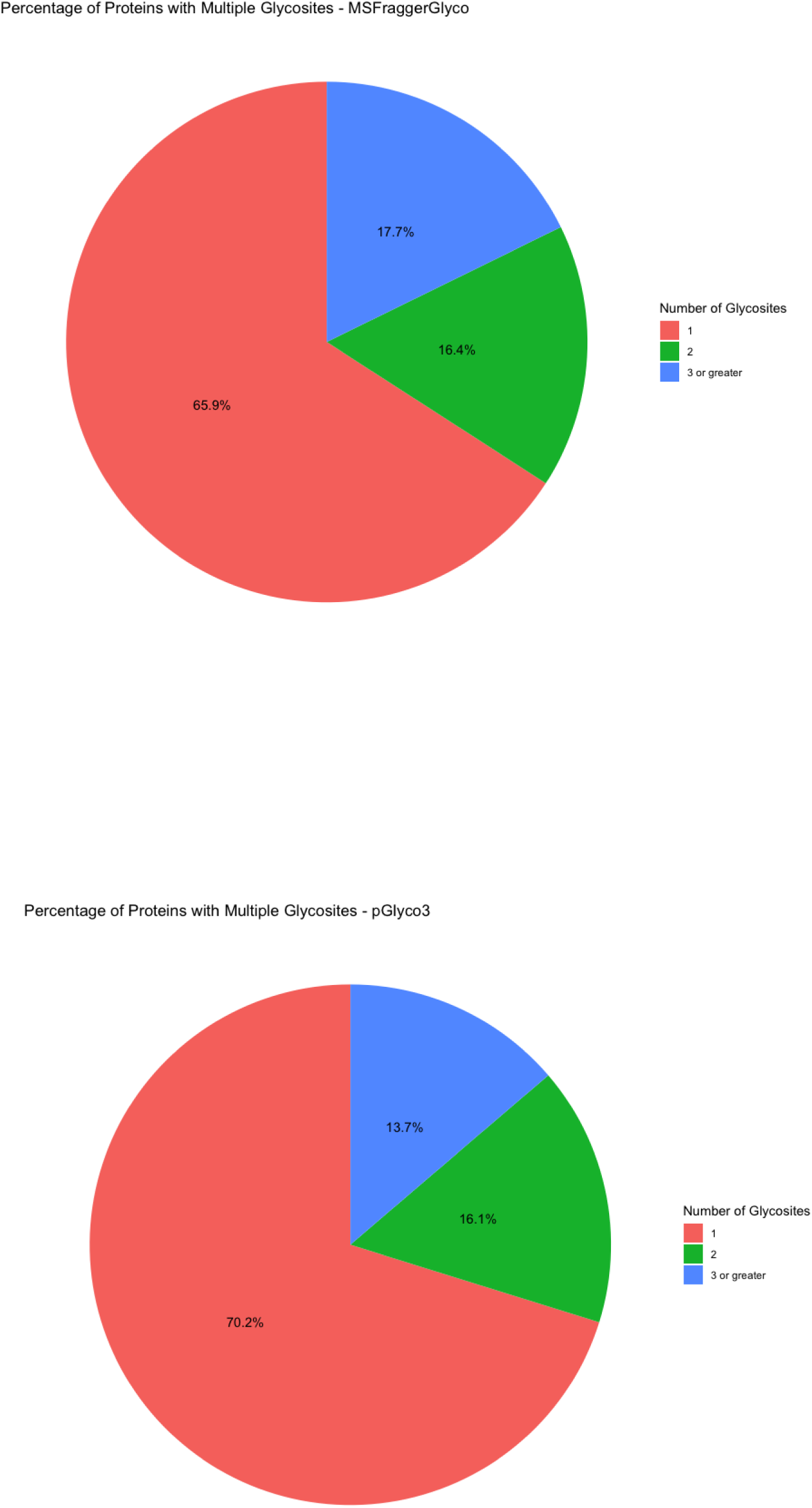
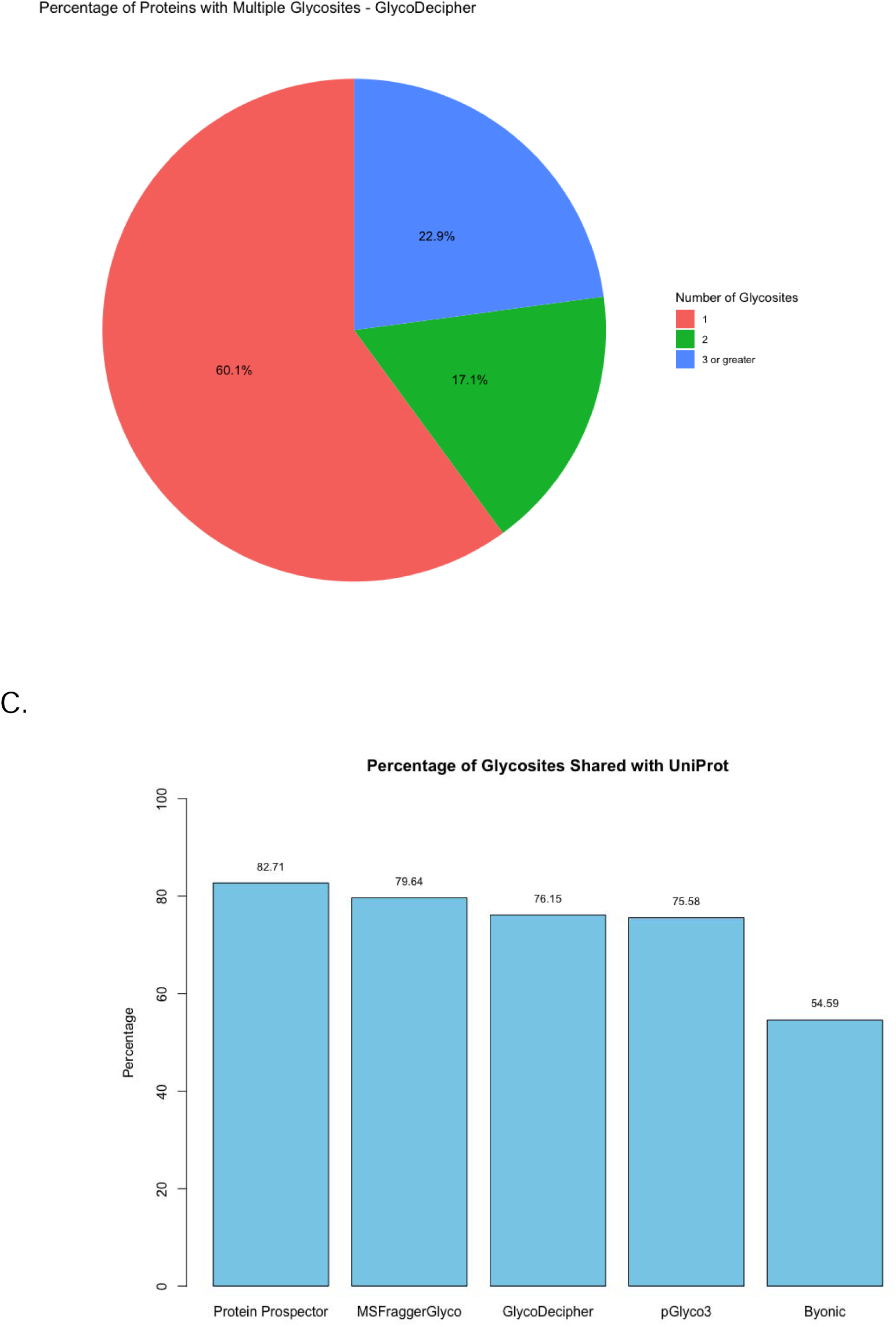
Comparison of the Unique Glycosites. (A) UpSet plot of the unique glycosites by software. (B) Pie Chart for the percentage of proteins with multiple glycosites (C) Percentage of glycosites also reported by UniProt.

There is a sizeable portion of proteins reported in each software to have multiple glycosites. Figure 5B shows that while most proteins have a single *N*-glycosite, somewhere between 24 to 40% have multiple. LRP1 is exceptional in that it has somewhere between 12 and 15 glycosites (**Supplementary Figure 3**), as has been observed in prior studies.(29) Overall, there is equivalent consensus on where glycosites are located (approximately 25% of glycosites reported by all softwares) as there is on what proteins are being glycosylated (approximately 24% of glycoproteins reported by all softwares).

A biologically relevant result from glycopeptide search software is the discovery of novel glycosites. Hence, glycosites for each UniProtID identified were compared to the glycosites discovered by the different software (Figure 5C). Not all glycosites reported in UniProt were expected in the results, but our goal was to understand what percent of those identified in our data are also reported in UniProt as a proxy for reliability. Byonic had the lowest agreement with UniProt with 54% of the sites identified also reported by UniProt. While it is possible to conclude from this that Byonic is more sensitive than competitors, a more likely conclusion given its consistently inflated numbers is that it is reporting more spurious results. For reference, Protein Prospector, which identified the most glycopeptide spectra and the fewest glycosites, had the highest agreement with 82.71% of its sites also being reported in UniProt.

### Comparison of Glycans

Overall, the software identified 99 distinct glycans (Figure 6A). Of these, 25 were common to all software. GlycoDecipher exclusively identified 11. This was expected because GlycoDecipher performs *de novo* “monosaccharide stepping” to report glycans not within the search space of other software. After looking for glycopeptides with glycans provided in its database, GlycoDecipher tries to identify more glycans by allowing for modifications on the monosaccharides to subsequently identify the additional peak/s in a Y-ion series. Most results from GlycoDecipher were identified in this manner but could be translated into defined glycan compositions (**Table 1**). In most cases, the additional mass was the result of a misassigned monoisotopic peak that, when corrected, led to a glycan assignment that was within the glycan database considered by other software. However, there were some glycans identified that were not identified in the glycomic analysis, notably those containing phosphate groups. Standards for these glycans were not available for glycomic analysis. These results suggest that GlycoDecipher’s *de novo* approach could be useful but is greatly hampered by its reliability.

**Figure 6:**
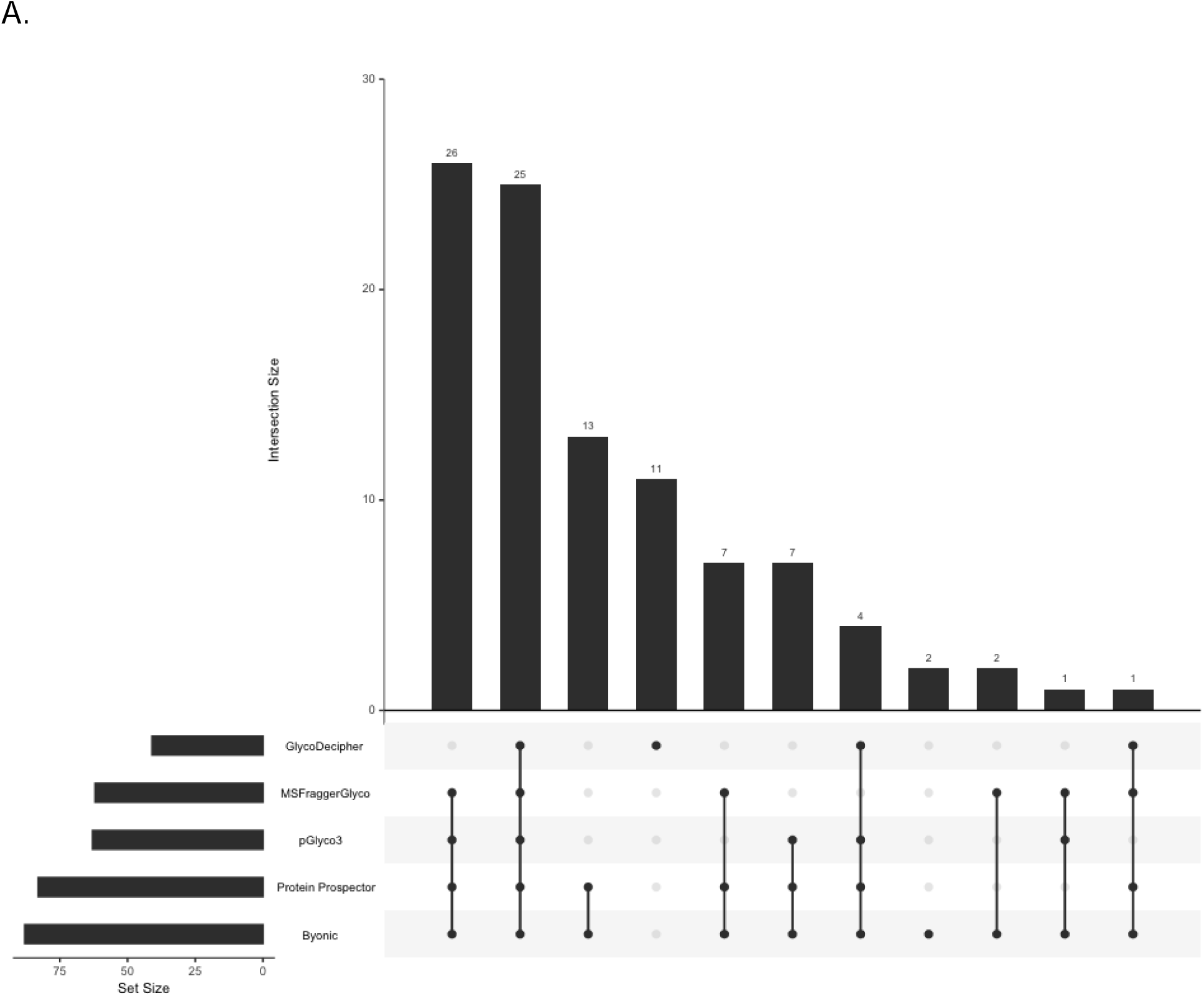
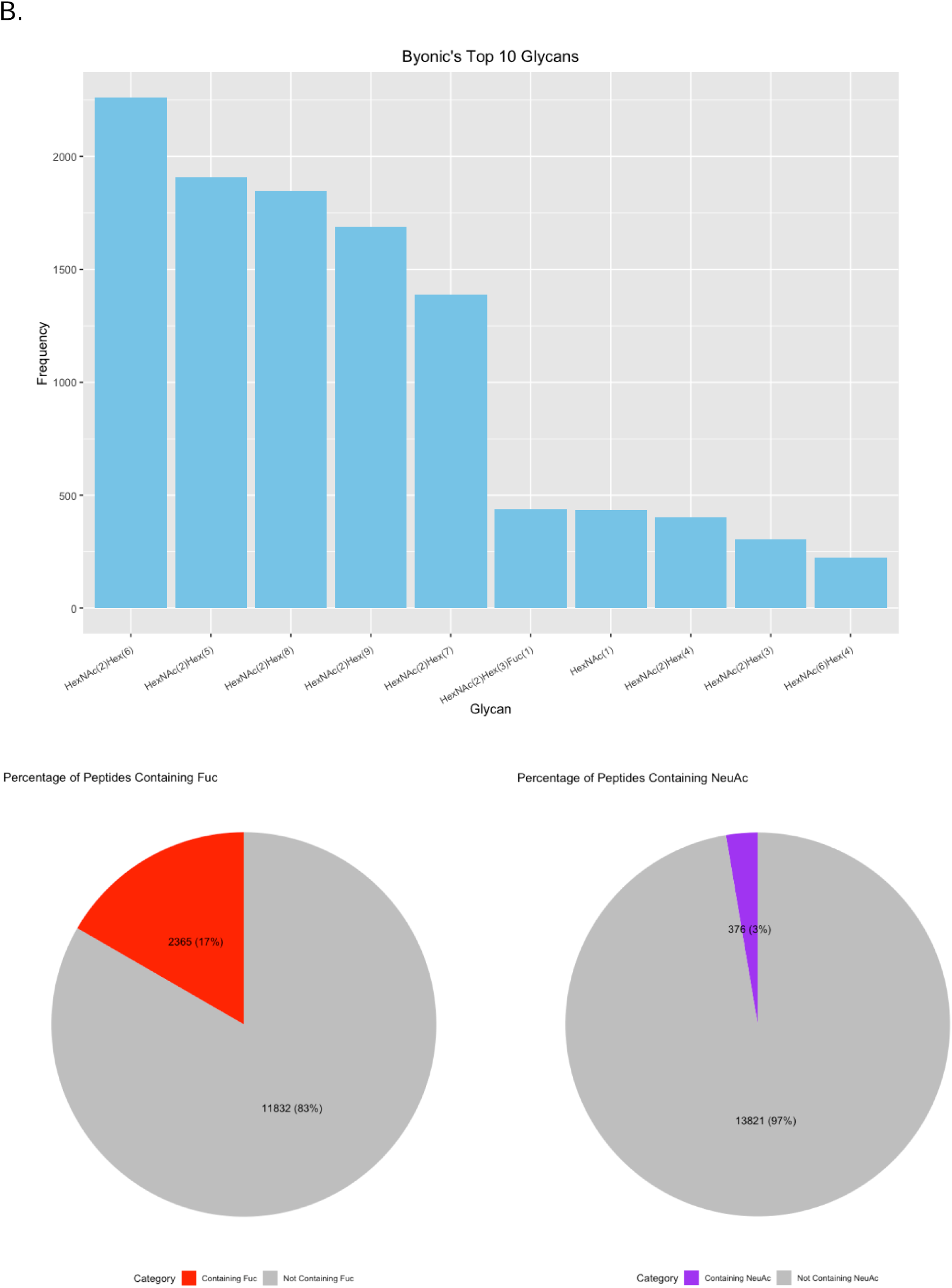
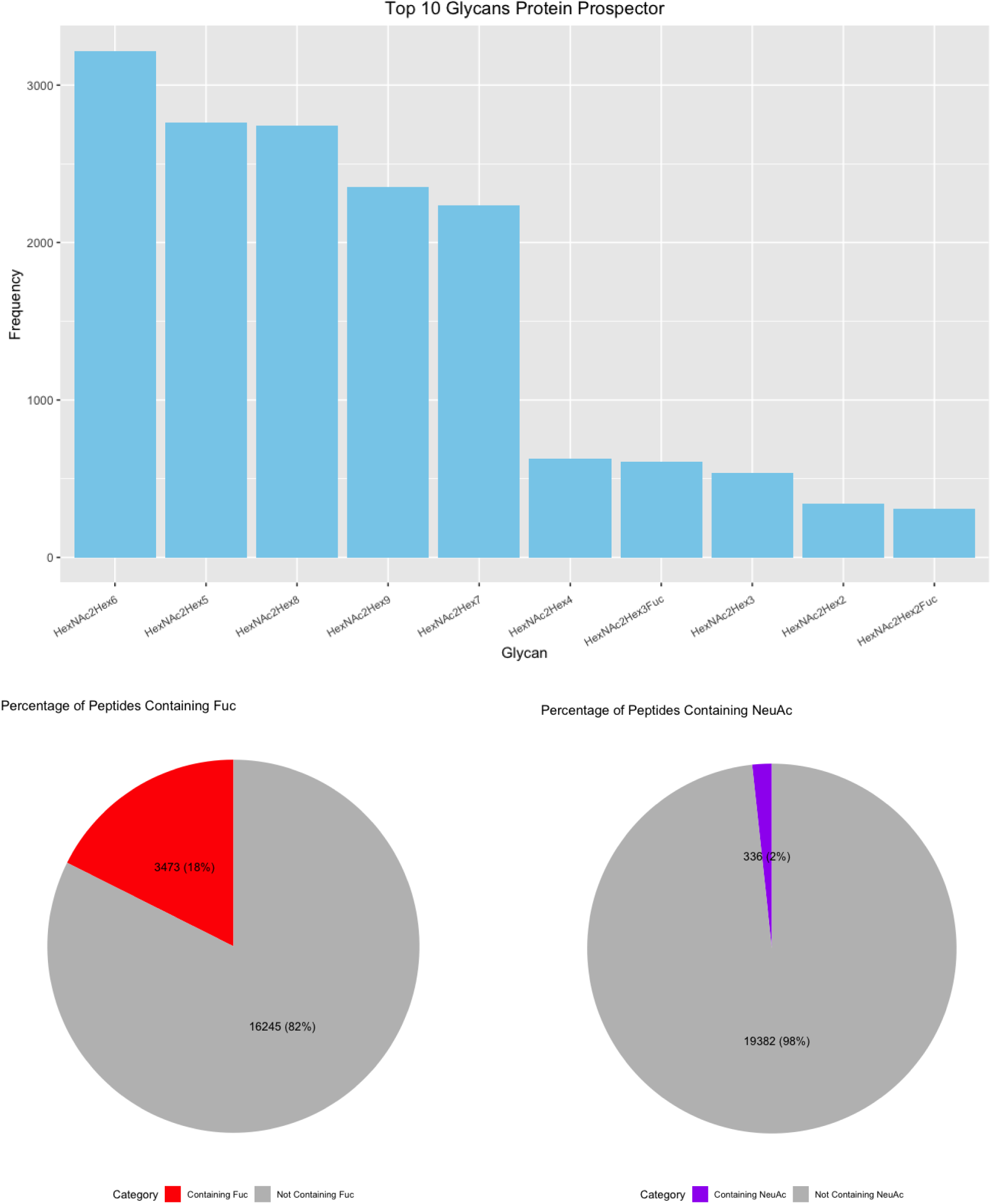
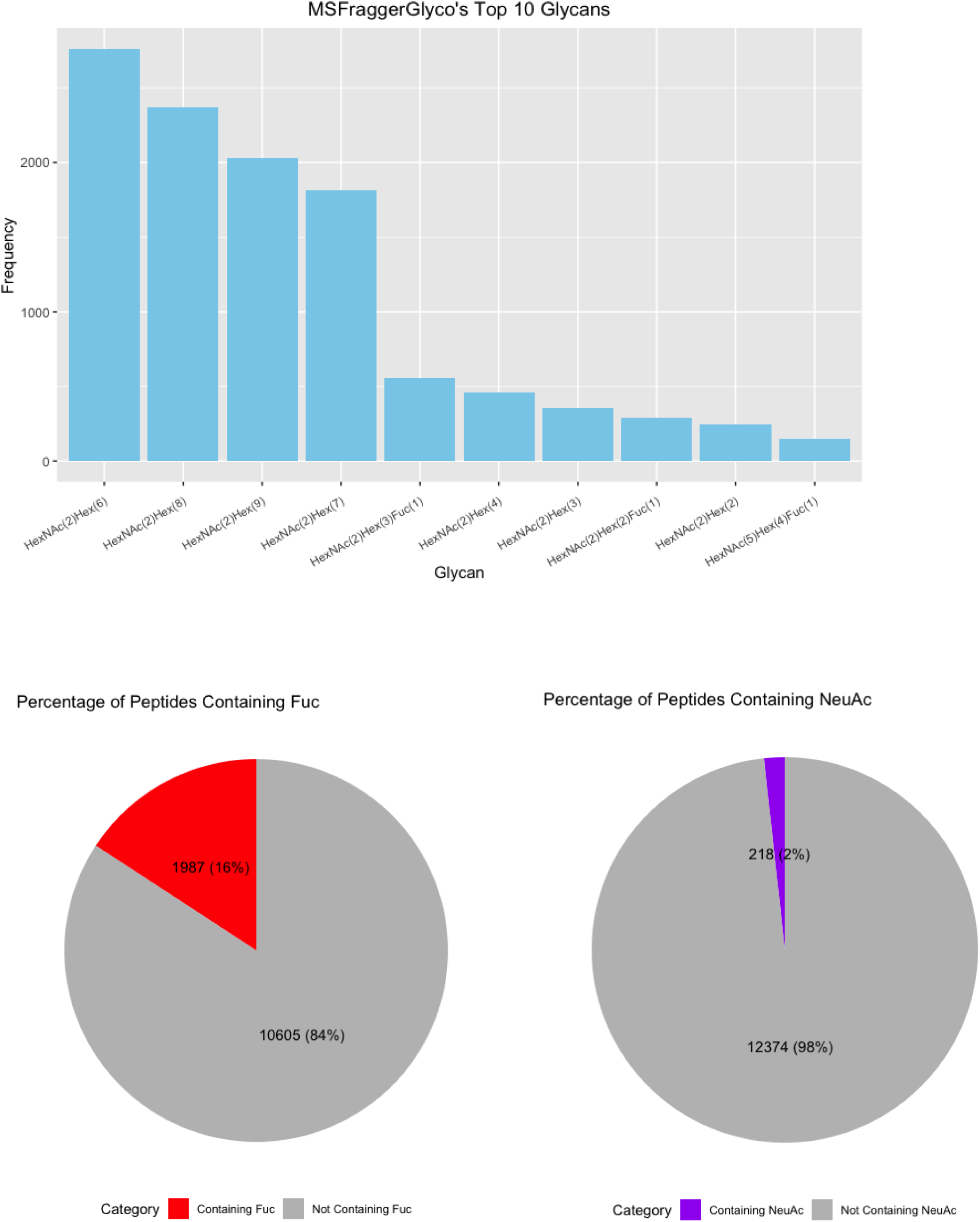
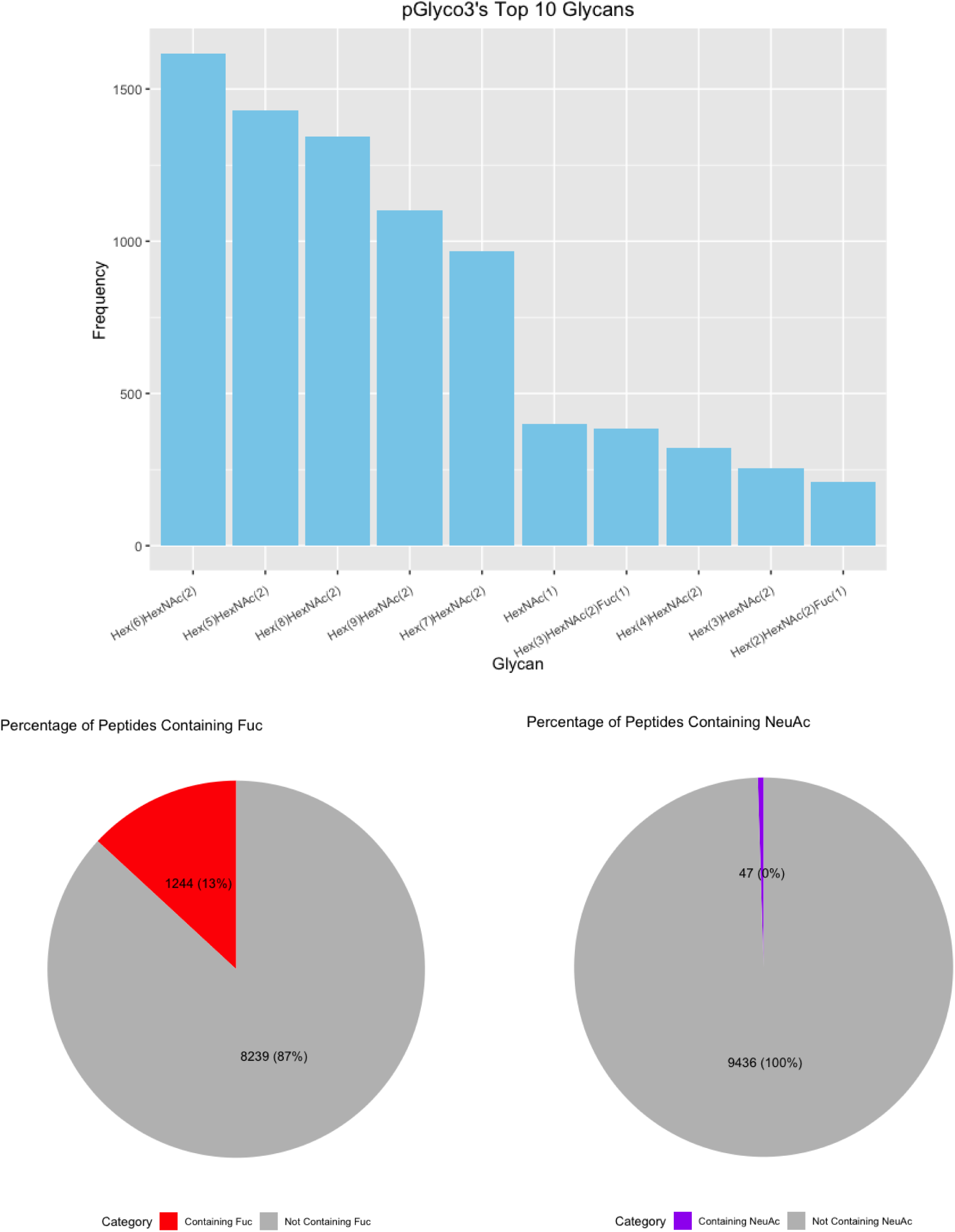
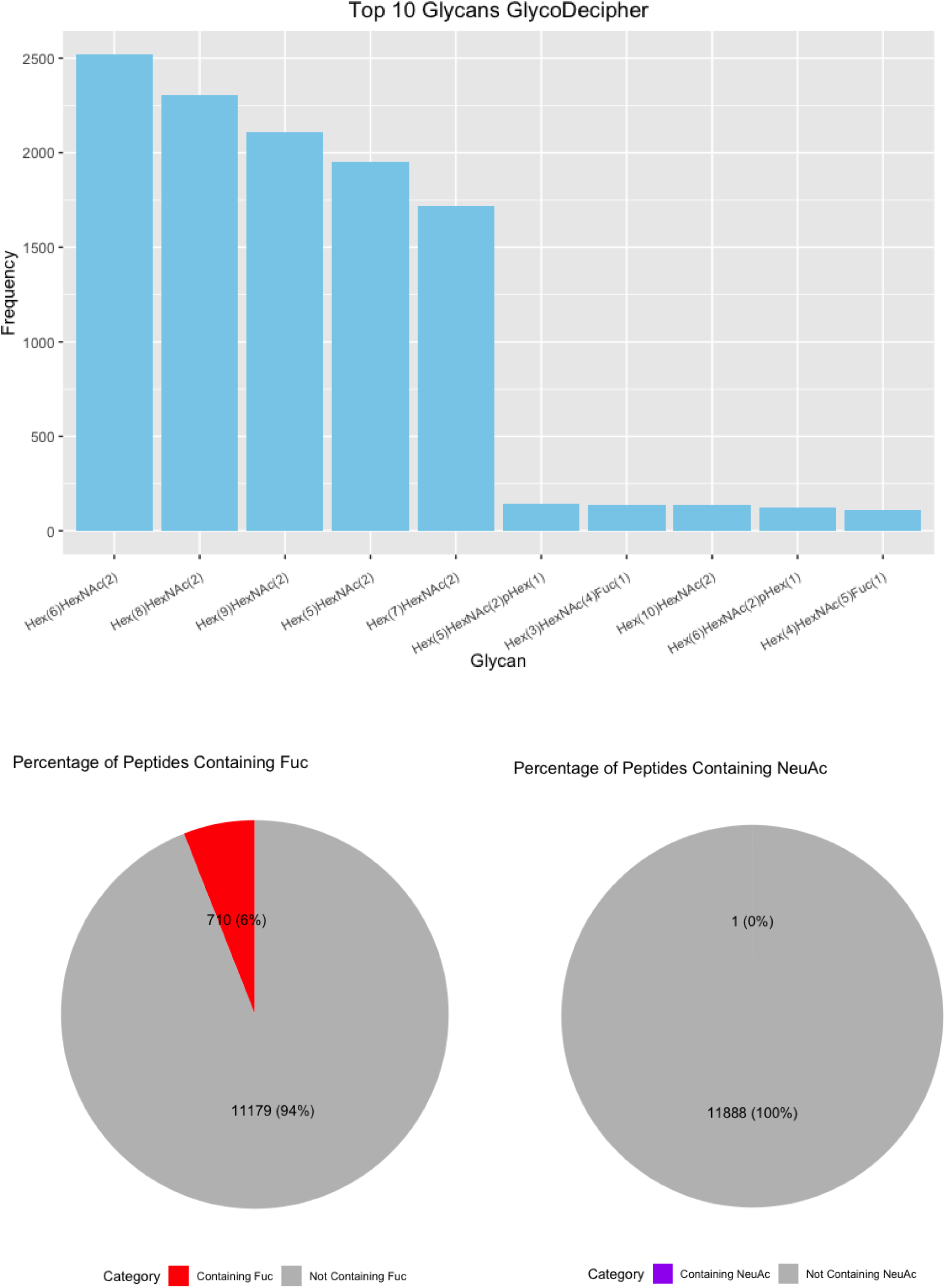
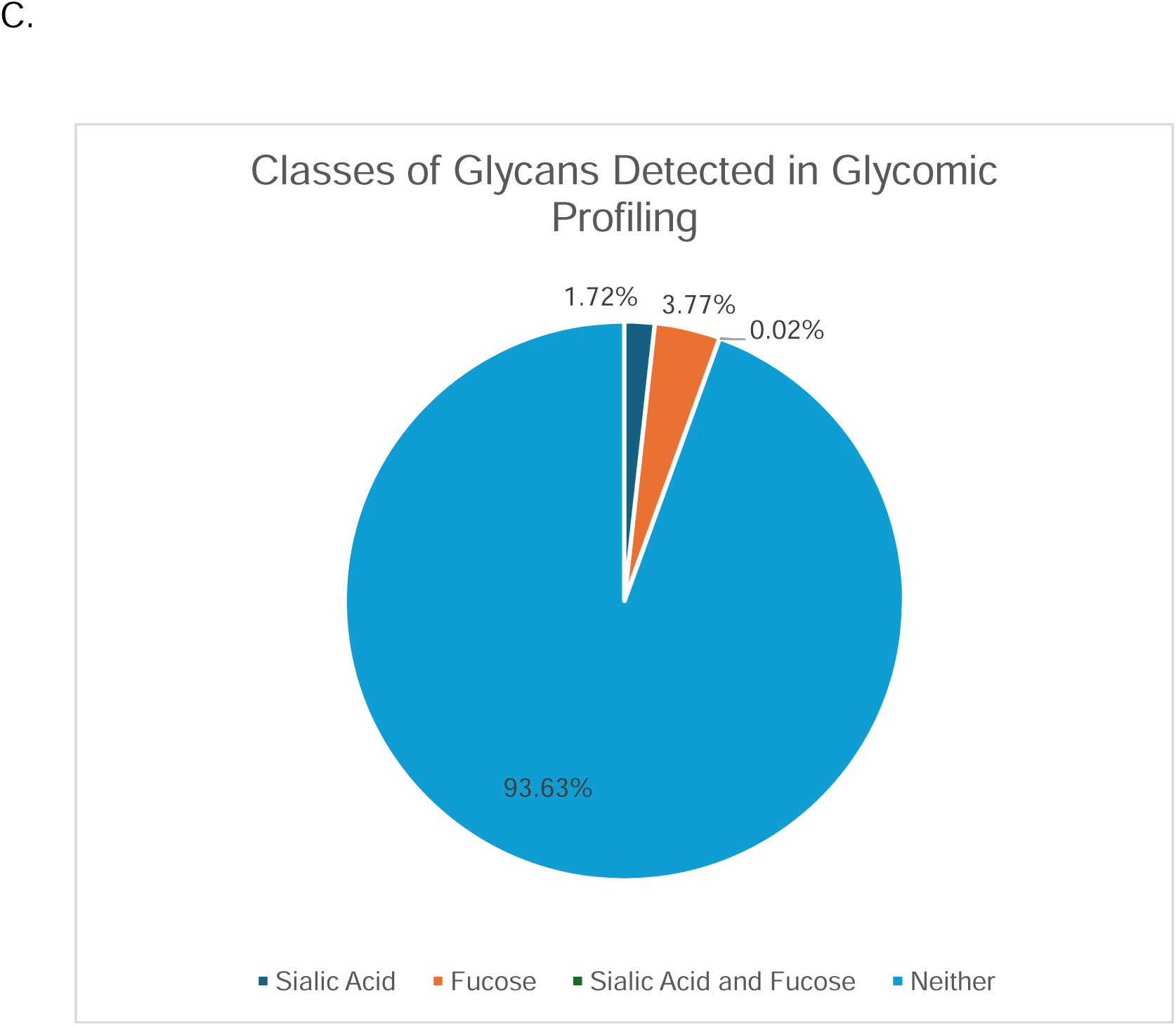
Comparison of the Unique Glycans. (A) Upset Plot of the glycans identified by each software. (B) Top 10 glycans identified by each software with pie charts displaying the percent of fucosylated and sialylated peptides. (C) Glycans from glycomic profiling split by features such as fucosylation and sialylation.

**Table 1:** Glycan Misassignments from GlycoDecipher. “Moiety Mass” is the mass of the nontraditional monosaccharide reported by GlycoDecipher. “Corresponds to:” shows our assessment of what the mass corresponds to based on the value and the spectra. “Operation” displays the function applied to correct for the misassignment. Highlighted rows are major structural errors. **Bold** values are examples of uncommon monosaccharides, such as a phosphorylated hexose.

| **Moiety Mass** | **Corresponds to:** | **Operation** |
| --- | --- | --- |
| 178.06 | Hexose + Oxygen | Add 2 Hexose, Subtract Fucose |
| 178.07 | Hexose + Oxygen | Add 2 Hexose, Subtract Fucose |
| 178.08 | Hexose + Oxygen | Add 2 Hexose, Subtract Fucose |
| 179.07 | Hexose + Oxygen, monoisotopic peak misassignment | Add 2 Hexose, Subtract Fucose |
| 179.08 | Hexose + Oxygen, monoisotopic peak misassignment | Add 2 Hexose, Subtract Fucose |
| 184.06 | Hexose + Na | Add Hexose |
| 187.11 | HexNAc - Oxygen | Add HexNAc, Add Fucose, Subtract Hexose |
| 187.12 | HexNAc - Oxygen | Add HexNAc, Add Fucose, Subtract Hexose |
| 188.12 | HexNAc - Oxygen, monoisotopic peak misassignment | Add HexNAc, Add Fucose, Subtract Hexose |
| **201.02** | **2 Hexoses + Phosphate** | **Subtract HexNAc, Add 2 Hexose, Add Phosphate** |
| **201.03** | **2 Hexoses + Phosphate** | **Subtract HexNAc, Add 2 Hexose, Add Phosphate** |
| 203.1 | HexNAc | Add HexNAc |
| 203.11 | HexNAc | Add HexNAc |
| 203.12 | HexNAc | Add HexNAc |
| 204.11 | HexNAc, monoisotopic peak misassignment | Add HexNAc |
| **241.08** | **Hexose + Phosphate** | **Add Hexose, Add Phosphate** |
| **242.04** | **Hexose + Phosphate** | **Add Hexose, Add Phosphate** |
| **242.05** | **Hexose + Phosphate** | **Add Hexose, Add Phosphate** |
| **243.04** | **Hexose + Phosphate** | **Add Hexose, Add Phosphate** |
| **251.11** | **Hexose + 2 Fucose** | **Add Hexose, Add 2 Fucose, Subtract HexNAc** |
| 267.11 | HexNAc - Oxygen + Phosphate | Add HexNAc, Add Fucose, Subtract Hexose, Add Phosphate |
| **280.99** | **5 Hexoses + Phosphate** | **Subtract 3 HexNAc, Add 5 Hexoses, Add Phosphate** |
| **283.07** | **HexNAc + Phosphate** | **Add HexNAc, Add Phosphate** |
| 307.12 | 2 Hexoses - Oxygen | Add Hexose, Add Fucose |
| 308.13 | 2 Hexoses - Oxygen | Add Hexose, Add Fucose |
| 308.14 | 2 Hexoses - Oxygen | Add Hexose, Add Fucose |
| 308.15 | 2 Hexoses - Oxygen | Add Hexose, Add Fucose |
| 309.15 | 2 Hexoses - Oxygen | Add Hexose, Add Fucose |
| 323.12 | 2 Hexoses, monoisotopic peak misassignment | Add 2 Hexoses |
| 323.13 | 2 Hexoses, monoisotopic peak misassignment | Add 2 Hexoses |
| 323.14 | 2 Hexoses, monoisotopic peak misassignment | Add 2 Hexoses |
| 324.11 | 2 Hexoses | Add 2 Hexoses |
| 324.12 | 2 Hexoses | Add 2 Hexoses |
| 324.13 | 2 Hexoses | Add 2 Hexoses |
| 324.14 | 2 Hexoses | Add 2 Hexoses |
| 324.15 | 2 Hexoses | Add 2 Hexoses |
| 324.16 | 2 Hexoses | Add 2 Hexoses |
| 325.12 | 2 Hexoses, monoisotopic peak misassignment | Add 2 Hexoses |
| 325.13 | 2 Hexoses, monoisotopic peak misassignment | Add 2 Hexoses |
| 325.14 | 2 Hexoses, monoisotopic peak misassignment | Add 2 Hexoses |
| 325.15 | 2 Hexoses, monoisotopic peak misassignment | Add 2 Hexoses |
| 326.13 | 2 Hexoses, monoisotopic peak misassignment | Add 2 Hexoses |
| 326.14 | 2 Hexoses, monoisotopic peak misassignment | Add 2 Hexoses |
| 327.15 | 2 Hexoses, monoisotopic peak misassignment | Add 2 Hexoses |
| 341.16 | 2 Hexoses + Ammonia | Add 2 Hexoses |

Finally, we investigated the frequency of each glycan reported. HexNAc_2_Hex_6_ was consistently the most identified glycan (Figure 6B). Interestingly, it was not the most abundant glycan from glycomic profiling, which was HexNAc_2_Hex_8_ (**Supplementary Table 2**). The glycoproteomic results were not evaluated for peak intensity, so this difference presumably indicates that HexNAc_2_Hex_6_ glycoforms, although more common on proteins, were typically of lower abundance than HexNAc_2_Hex_8_. An alternate, less likely explanation could also include a small degree of high mannose glycan degradation during sample prep or in the gas-phase. Regardless, HexNAc_2_Hex_8_ was either the second or third most frequent glycan in glycoproteomic searches (Figure 6B). HexNAc_2_Hex_9_ was the second most abundant glycan by glycomic profiling and was the third or fourth most frequent glycan depending on the software. Taken together, all software identified the most common glycans but differed in their detection frequency. Software also differed in the low abundance glycans detected.

We also investigated the percentage fucosylation and sialylation. Very little sialylation was found in this data: 1% or less of IDs by all software. These results matched that of the glycomic profiling (Figure 6C). Fucosylation was more common with up to 20% of glycopeptides reported by a given software. Interestingly, this differed from the glycomic profiling results, where fucosylated glycans were only 3.77% of the glycans identified. A common error in data analysis is to assign two fucose where there is only a sialic acid because two fucose (292.1158 Da) and a sialic acid (291.0954) are close in mass. **Supplementary Figure 4** displays the percent of peptides with multiple fucose. Only up to 2% of the glycopeptides in any software contained two fucose, which makes it an uncommon occurrence. The high frequency of fucosylation on glycopeptides relative to the low abundance of fucosylated glycan by glycomic profiling is not due to misassignment. These data suggest that counting the frequency of a glycan on peptides is a poor proxy for its relative abundance; peak intensities need to be taken into consideration.

### Usability of Glycoproteomic Software

Usability of a software greatly influences its longevity with users. A computer with ample RAM is critical for MSFraggerGlyco. All 24 RAW files could not be searched with the 16GB of RAM available on a high-end laptop or typical desktop computer. Using a computer with 1 TB of RAM solved this problem. We did not evaluate the minimum RAM required for MSFragger to complete this search. Provided that there is appropriate RAM, MSFraggerGlyco was the fastest software, completing its search in 91.4 minutes (**Supplementary Figure 5**). Byonic was the slowest to complete its search at 364.1 minutes.

The ability to recalibrate the data is useful for a robust glycoproteomic analytical tool. Due to the high mass accuracy of modern mass spectrometers a small systematic error in calibration can impact search performance. The data in this study had a mass error of on average 8 ppm. Every software except Byonic had either a wider default precursor tolerance or automatically adjusted for this systematic error. To get comparable results, Byonic required manual adjustment to a 20 ppm precursor tolerance (**Supplementary Figure 6**). Some software, such as MSFraggerGlyco and pGlyco3, have a larger precursor tolerance and did not require adjustment. Protein Prospector and GlycoDecipher had narrower tolerances but could adjust for the systematic error.

## Discussion

In this work, we complete a head-to-head comparison of five contemporary glycoproteomic analytical software: Byonic, Protein Prospector, MSFraggerGlyco, pGlyco3, and GlycoDecipher. These five were chosen because of their blend of features that have different advantages. We note that this study focused on *N*-glycoproteins and did not include tools or analyses more tailored for *O*-glycoproteins.(19,30,31) Byonic searches for a glycan as a variable modification on the peptide. Protein Prospector and MSFraggerGlyco determine the difference in mass between the precursor and the unfragmented peptide backbone and use the mass offset to define the mass and composition of the glycan. pGlyco3 and GlycoDecipher filter spectra for those that contain sufficient Y-ions, which are unfragmented peptides with fragmented glycans, and construct the glycan from these fragments. Additionally, MSFraggerGlyco, pGlyco3, and GlycoDecipher all calculate a false discovery rate for the glycan to generate an estimate of confidence in reported results.

Based on our results, a single winner is not evident; but there are important lessons. One clear finding is that although Byonic was a gold standard in this field and has enabled the development of other tools, it appears to produce results of lower reliability than modern alternatives. It reports the most unique proteins and glycosites but does not identify the most glycopeptide spectra. By spot-checking, we show that some of these are misassignments. Additionally, almost 50% of the glycosites reported by Byonic could not be supported by UniProt. While Uniprot is in no way comprehensive, one can feel more confident in glycosite assignments that have been previously reported. It is possible that Byonic performs poorly because it attempts to identify all glycan and peptide ions at once and reports confidence based on scores for the presence of certain ions. In the example we show, Byonic may score noisy peaks very highly while ignoring critical information, such as the most abundant peaks in the spectra.

Glycoproteomic search engines should be selected based on the goals of the experiment. If one wants to identify the most glycopeptides as in an exploratory experiment, then mass offset approaches would be most favored. Protein Prospector was able to identify the most glycopeptides of any software. Among software calculating a glycan FDR, MSFraggerGlyco reported the most spectra. Whether the reported glycan FDR is accurate is open to question. Although Protein Prospector does not calculate a glycan FDR and instead uses a confidence score, it found the greatest number of unique combinations of protein, site, and glycan (PSG IDs). This did not sacrifice accuracy, at least with metrics used here for glycoprotein and glycosite assignments, since it still reported the least number of glycosites and maintained the highest agreement with UniProt.

If one wants to focus on the high-confidence structure of the glycan at a given site, pGlyco3 is a strong choice. Although it does not report the most glycopeptides, its Y-ion approach and glycan FDR calculation provide greater confidence in the assignment. Put differently, rather than inferring a glycan from its mass, it relies on direct evidence from peaks in the spectrum. Still, it has been reported to have a bias in assigning more fucosylated peptides although recent fixes seem to have solved this problem as other software reported more fucosylation than pGlyco3.(32) GlycoDecipher’s *de novo* monosaccharide stepping suffers from problems with accurate glycan assignment. Manual inspection revealed that the overwhelming majority of glycans assigned as novel could be explained by glycans within the supplied glycan database. Despite that, it is possible to correct the results; so there is a clear path forward for this software to improve.

There is clear benefit to users trying more than one software for a given dataset. Although all software agreed on a core set of results, the next largest subsets were the unique results of a single software. At least two software, excluding Byonic, agreed on over half the results. It may prove useful to use one mass offset approach and one Y-ion approach. The mass offset approach will identify the most candidate glycopeptides. The Y-ion approach will provide those glycopeptides with strong Y-ion coverage, which is important for confident glycan assignment. Taken together, these approaches offer complementary information from the peptide and the glycan.

Finally, the use of glycomic profiling to generate a standardized glycan database provided greater confidence in assignments, enabled a rigorous comparison, and illustrated what features of a glycopeptide software are useful. There are currently few options to validate glycopeptide assignments. One can spot check for a specific protein by exo- or endoglycosidase digestion followed by Western blotting, but this is low throughput. One can use lectin microarrays, but these are custom and costly. Glycomic profiling is one way to corroborate the glycan assignments and to limit results to only those glycans which can be identified by other means. However, the GlycoDecipher *de novo* results did uncover that glycans outside of the database are present even after manual correction, so it is important that standards for as many glycans as possible are available during the glycomic analysis. Here, we used the glycomic results to ensure that software were operating in similar search spaces. While this may not benefit every researcher, it allowed a fair comparison of software and was instrumental to discovering the sensitivity of the mass offset approach.

Converting the glycomic results into a database for each glycoproteomic software was challenging. This is partly because some software use input formats that include glycan topology whereas others just use compositions; but even among software that use the same type of input, there are still formatting differences. For example, GlyTouCan provides a universal database for all glycans and can provide unique identifiers to different levels of resolution (monosaccharide composition, isomer composition, topology, and linkage), but using these identifiers would be extremely challenging for comparing software as each software reports different identifiers for the same assignment. A universal glycan database format would be critical to establishing better consistency between software and would eliminate a significant amount of work involved in reformatting databases.

## Acknowledgments

The authors would like to acknowledge Christophe Guilbert for setting up and managing the hardware on which all searches were run. Figure 1 was made with BioRender.com. RAW files, search engine output files, fasta files, glycan databases, R scripts, Python scripts, and bash scripts used in this work have been deposited to the ProteomeXchange Consortium via MassIVE partner repository with the dataset identifier MSV000095432 or PXD054193.

## Author Contribution

R.A.H. designed and performed the data acquisition and analysis. L.E.P. performed glycomic profiling. N.M.R. provided guidance for sample preparation and data acquisition. R.A.H. and R.J.C. wrote the manuscript. R.J.C. supervised all data analysis.

