## Supplementary Materials for "“Comparative Analysis of Glycoproteomic Software Using a Tailored Glycan Database”"

**PDF includes:**

Supplementary Tables 1 to 2

Supplementary Figures 1 to 6

**Supplementary Table 1: Functional Enrichment Results for All Peptides Identified by MaxQuant.** Results were filtered by the UniProt Keywords category and sorted by FDR value.

| # background genes | # genes | description | FDR value | p-value |
|---|---|---|---|---|
| 8122 | 4237 | Phosphoprotein | 1.06E-298 | 1.58E-301 |
| 3362 | 2476 | Acetylation | 2.24E-278 | 6.66E-281 |
| 5095 | 2419 | Cytoplasm | 1.03E-80 | 4.61E-83 |
| 1717 | 1072 | Isopeptide bond | 2.68E-70 | 1.60E-72 |
| 2399 | 1344 | Ubl conjugation | 3.46E-68 | 2.57E-70 |
| 5278 | 2413 | Nucleus | 4.84E-68 | 4.32E-70 |
| 10313 | 3926 | Alternative splicing | 8.74E-48 | 9.10E-50 |
| 1181 | 719 | Mitochondrion | 1.61E-42 | 1.91E-44 |
| 686 | 481 | RNA-binding | 1.90E-38 | 2.55E-40 |
| 617 | 433 | Protein transport | 1.55E-34 | 2.31E-36 |
| 973 | 583 | Methylation | 1.21E-32 | 1.99E-34 |
| 545 | 382 | Transit peptide | 2.67E-30 | 4.77E-32 |
| 278 | 236 | Ribonucleoprotein | 3.43E-26 | 6.64E-28 |
| 349 | 271 | mRNA processing | 5.65E-26 | 1.18E-27 |
| 1776 | 859 | Nucleotide-binding | 1.74E-24 | 3.89E-26 |
| 274 | 225 | mRNA splicing | 1.02E-23 | 2.43E-25 |
| 1379 | 673 | ATP-binding | 1.69E-19 | 4.28E-21 |
| 175 | 154 | Ribosomal protein | 4.44E-18 | 1.19E-19 |
| 540 | 322 | Host-virus interaction | 1.71E-17 | 4.85E-19 |
| 2166 | 931 | Coiled coil | 5.35E-15 | 1.59E-16 |
| 138 | 121 | Spliceosome | 3.63E-14 | 1.13E-15 |
| 349 | 221 | Neurodegeneration | 5.91E-14 | 1.93E-15 |
| 292 | 195 | Chromatin regulator | 6.18E-14 | 2.16E-15 |
| 1235 | 578 | Cytoskeleton | 6.18E-14 | 2.12E-15 |
| 651 | 345 | Cell cycle | 2.81E-13 | 1.04E-14 |
| 128 | 108 | Protein biosynthesis | 5.35E-12 | 2.07E-13 |
| 197 | 142 | Chaperone | 5.94E-12 | 2.39E-13 |
| 384 | 225 | Cell division | 9.84E-12 | 4.10E-13 |
| 157 | 121 | Primary mitochondrial disease | 1.39E-11 | 6.00E-13 |
| 95 | 87 | rRNA processing | 5.20E-11 | 2.32E-12 |
| 1978 | 816 | Transport | 3.80E-10 | 1.76E-11 |
| 282 | 172 | WD repeat | 4.91E-10 | 2.34E-11 |
| 433 | 236 | Chromosome | 5.05E-10 | 2.48E-11 |

| | | | | |
|---|---|---|---|---|
| 297 | 175 | Mitochondrion inner membrane | 2.35E-09 | 1.19E-10 |
| 141 | 103 | Helicase | 4.98E-09 | 2.59E-10 |
| 275 | 163 | Mitosis | 6.91E-09 | 3.70E-10 |
| 83 | 73 | Ribosome biogenesis | 8.06E-09 | 4.44E-10 |
| 105 | 84 | mRNA transport | 9.93E-09 | 5.62E-10 |
| 386 | 208 | DNA damage | 1.33E-08 | 7.71E-10 |
| 1820 | 736 | Transferase | 5.37E-08 | 3.20E-09 |
| 325 | 178 | DNA repair | 8.54E-08 | 5.24E-09 |
| 1612 | 660 | Hydrolase | 8.54E-08 | 5.21E-09 |
| 91 | 71 | ER-Golgi transport | 3.69E-07 | 2.36E-08 |
| 695 | 316 | Ubl conjugation pathway | 8.98E-07 | 5.88E-08 |
| 131 | 88 | Ligase | 1.19E-06 | 7.97E-08 |
| 568 | 265 | Endosome | 1.95E-06 | 1.33E-07 |
| 86 | 65 | Translocation | 2.91E-06 | 2.04E-07 |
| 138 | 88 | Centromere | 5.79E-06 | 4.14E-07 |
| 108 | 74 | Neuropathy | 6.59E-06 | 4.81E-07 |
| 52 | 46 | Proteasome | 8.41E-06 | 6.26E-07 |
| 98 | 68 | DNA replication | 1.31E-05 | 9.95E-07 |
| 1168 | 476 | Endoplasmic reticulum | 1.64E-05 | 1.27E-06 |
| 117 | 76 | Translation regulation | 1.89E-05 | 1.49E-06 |
| 100 | 68 | Kinetochore | 2.06E-05 | 1.66E-06 |
| 53 | 45 | Nuclear pore complex | 2.15E-05 | 1.76E-06 |
| 107 | 71 | Electron transport | 2.25E-05 | 1.87E-06 |
| 271 | 139 | Actin-binding | 4.14E-05 | 3.56E-06 |
| 38 | 36 | Aminoacyl-tRNA synthetase | 4.14E-05 | 3.58E-06 |
| 63 | 49 | Respiratory chain | 4.14E-05 | 3.51E-06 |
| 578 | 257 | Magnesium | 4.20E-05 | 3.75E-06 |
| 280 | 142 | Microtubule | 4.88E-05 | 4.43E-06 |
| 570 | 253 | Oxidoreductase | 5.33E-05 | 4.91E-06 |
| 611 | 266 | Repressor | 9.29E-05 | 8.71E-06 |
| 90 | 60 | tRNA processing | 1.10E-04 | 1.02E-05 |
| 47 | 39 | Initiation factor | 1.20E-04 | 1.21E-05 |
| 131 | 78 | Isomerase | 1.20E-04 | 1.16E-05 |
| 125 | 74 | Mitochondrion outer membrane | 2.20E-04 | 2.20E-05 |
| 51 | 39 | Redox-active center | 4.30E-04 | 4.34E-05 |

| | | | | |
|---|---|---|---|---|
| 312 | 145 | Chromosomal rearrangement | 9.60E-04 | 9.90E-05 |
| 55 | 39 | Charcot-Marie-Tooth disease | 0.0013 | 1.40E-04 |
| 2369 | 860 | Transcription | 0.0013 | 1.40E-04 |
| 37 | 30 | DNA-directed RNA polymerase | 0.0015 | 1.60E-04 |
| 879 | 349 | Golgi apparatus | 0.0015 | 1.60E-04 |
| 595 | 248 | Cytoplasmic vesicle | 0.0015 | 1.60E-04 |
| 342 | 154 | GTP-binding | 0.0018 | 2.00E-04 |
| 163 | 83 | TPR repeat | 0.0031 | 3.50E-04 |
| 156 | 80 | Autophagy | 0.0033 | 3.80E-04 |
| 34 | 27 | Amyotrophic lateral sclerosis | 0.0035 | 4.10E-04 |
| 30 | 25 | Nonsense-mediated mRNA decay | 0.0035 | 4.10E-04 |
| 534 | 221 | Apoptosis | 0.0039 | 4.70E-04 |
| 27 | 23 | tRNA-binding | 0.0046 | 5.60E-04 |
| 634 | 255 | Kinase | 0.0055 | 6.70E-04 |
| 186 | 90 | Methyltransferase | 0.0059 | 7.30E-04 |
| 212 | 99 | Epilepsy | 0.0082 | 0.001 |
| 25 | 21 | Amino-acid biosynthesis | 0.0083 | 0.0011 |
| 128 | 66 | Endocytosis | 0.0083 | 0.001 |
| 35 | 26 | Leukodystrophy | 0.0083 | 0.001 |
| 55 | 35 | RNA-mediated gene silencing | 0.0092 | 0.0012 |
| 175 | 84 | S-adenosyl-L-methionine | 0.0097 | 0.0013 |
| 390 | 164 | Serine/threonine-protein kinase | 0.0104 | 0.0014 |
| 26 | 21 | rRNA-binding | 0.0112 | 0.0015 |
| 35 | 25 | Rotamase | 0.0139 | 0.0019 |
| 120 | 61 | Flavoprotein | 0.0147 | 0.002 |
| 3702 | 1274 | Metal-binding | 0.0156 | 0.0022 |
| 51 | 32 | Coated pit | 0.0158 | 0.0022 |
| 251 | 111 | Lipid-binding | 0.0158 | 0.0022 |
| 58 | 35 | Hereditary spastic paraplegia | 0.0162 | 0.0023 |
| 2299 | 811 | Transcription regulation | 0.0174 | 0.0025 |
| 45 | 29 | Congenital disorder of glycosylation | 0.0181 | 0.0027 |
| 66 | 38 | Citrullination | 0.0192 | 0.0029 |
| 39 | 26 | Bromodomain | 0.0206 | 0.0031 |
| 103 | 53 | Peroxisome | 0.0209 | 0.0032 |

| | | | | |
|---|---|---|---|---|
| 23 | 18 | Tricarboxylic acid cycle | 0.0265 | 0.0041 |
| 19 | 16 | Diamond-Blackfan anemia | 0.0265 | 0.0041 |
| 108 | 54 | FAD | 0.0293 | 0.0046 |
| 59 | 34 | Iron-sulfur | 0.0293 | 0.0046 |
| 50 | 30 | Exonuclease | 0.0307 | 0.0049 |
| 184 | 83 | NAD | 0.0311 | 0.005 |
| 74 | 40 | Multifunctional enzyme | 0.0317 | 0.0051 |
| 684 | 261 | Activator | 0.0323 | 0.0053 |
| 127 | 61 | Nuclease | 0.0325 | 0.0054 |
| 117 | 57 | Alternative initiation | 0.0333 | 0.0056 |
| 164 | 75 | Guanine-nucleotide releasing factor | 0.0338 | 0.0057 |
| 167 | 76 | Prenylation | 0.0349 | 0.0059 |
| 47 | 28 | Allosteric enzyme | 0.0402 | 0.0069 |
| 190 | 84 | GTPase activation | 0.041 | 0.0071 |
| 19 | 15 | One-carbon metabolism | 0.0458 | 0.008 |
| 17 | 14 | Mitochondrion nucleoid | 0.0458 | 0.008 |
| 170 | 76 | NADP | 0.0463 | 0.0082 |

**Supplementary Table 2**: **Glycans Identified in Glycomic Profiling.**

| GlyTouCan | Composition | Structure | Percent Abundance |
|---|---|---|---|
| G40702WU | (Hex)5 + (Man)3(GlcNAc)2 | 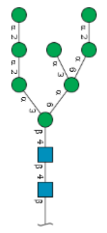 | 36.58% |
| G60230HH | (Hex)6 + (Man)3(GlcNAc)2 | 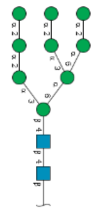 | 25.65% |
| G55220VL | (Hex)2 + (Man)3(GlcNAc)2 | 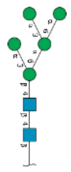 | 8.51% |
| G82933ZI | (HexNAc)4 (Deoxyhexose)1 + (Man)3(GlcNAc)2 | 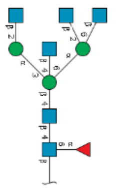 | 7.01% |
| G89993FE | (Hex)1 (HexNAc)2 + (Man)3(GlcNAc)2 | 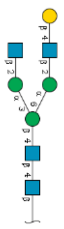 | 3.10% |
| G22573RC | (Man)2 (GlcNAc)2 | 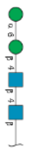 | 2.37% |

| | | | |
|---|---|---|---|
| G69411IG | (Hex)1 (HexNAc)1 + (Man)3(GlcNAc)2 | 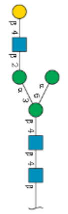 | 1.70% |
| G20030CU | (HexNAc)4 + (Man)3(GlcNAc)2 | 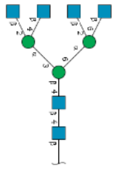 | 1.29% |
| G22768VO | (Man)3 (GlcNAc)2 | 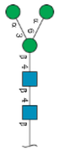 | 1.25% |
| G25520XG | (HexNAc)3 (Hex)2 (Deoxyhexose)1 + (Man)3(GlcNAc)2 | 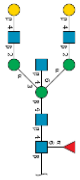 | 1.23% |
| G80966KZ | (Hex)3 + (Man)3(GlcNAc)2 | 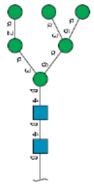 | 1.07% |
| G04216GB | (Hex)4 + (Man)3(GlcNAc)2 | 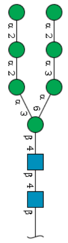 | 1.02% |

| | | | |
|---|---|---|---|
| G65890UA | (Hex)3 (HexNAc)1 (NeuAc)1 + (Man)3(GlcNAc)2 | 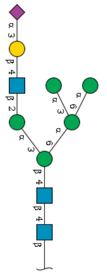 | 0.79% |
| G04576KS | (Hex)3 (HexNAc)1 (NeuAc)1 + (Man)3(GlcNAc)2 | 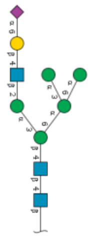 | 0.75% |
| G43677TG | (HexNAc)2 (Hex)1 + (Man)3(GlcNAc)2 | 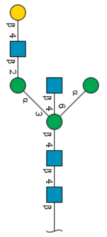 | 0.75% |
| G30588SI | (HexNAc)2 (Hex)2 + (Man)3(GlcNAc)2 | 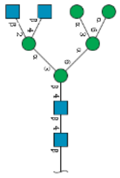 | 0.66% |
| G04414GO | (Hex)4 + (Man)3(GlcNAc)2 | 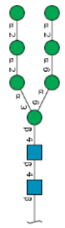 | 0.60% |
| G20966TZ | (Hex)2 (HexNAc)2 + (Man)3(GlcNAc)2 | 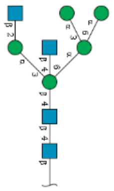 | 0.59% |

| | | | |
|---|---|---|---|
| G82942ZJ | (Hex)2 (HexNAc)1 (Deoxyhexose)1 + (Man)3(GlcNAc)2 | 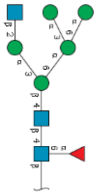 | 0.53% |
| G07617FP | (Hex)2 + (Man)3(GlcNAc)2 | 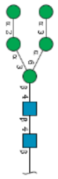 | 0.43% |
| G10219AA | (HexNAc)3 + (Man)3(GlcNAc)2 | 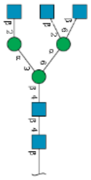 | 0.39% |
| G29011JC | (HexNAc)3 + (Man)3(GlcNAc)2 | 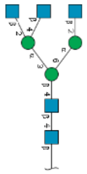 | 0.38% |
| G59471TH | (HexNAc)3 + (Man)3(GlcNAc)2 | 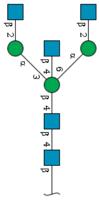 | 0.36% |
| G42227JK | (Man)3 (GlcNAc)2 | 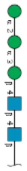 | 0.35% |
| G30159WR | (HexNAc)3 (Deoxyhexose)1 + (Man)3(GlcNAc)2 | 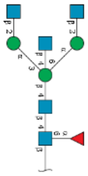 | 0.34% |

| | | | |
|---|---|---|---|
| G16828VN | (Hex)3 (HexNAc)1 + (Man)3(GlcNAc)2 | 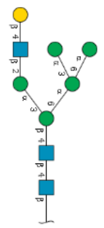 | 0.27% |
| G04675YX | (Hex)2 (HexNAc)2 (Deoxyhexose)1 + (Man)3(GlcNAc)2 | 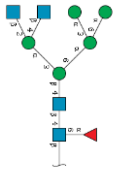 | 0.22% |
| G52934AK | (HexNAc)3 (Hex)2 + (Man)3(GlcNAc)2 | 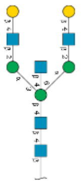 | 0.22% |
| G06374YY | (Hex)3 (HexNAc)1 (Deoxyhexose)1 + (Man)3(GlcNAc)2 | 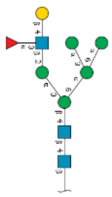 | 0.21% |
| G56014GC | (Hex)1 + (Man)3(GlcNAc)2 | 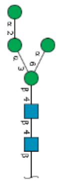 | 0.21% |
| G36191CD | (Hex)2 (HexNAc)2 + (Man)3(GlcNAc)2 | 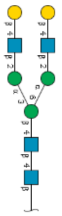 | 0.20% |
| G80858MF | (HexNAc)2 (Deoxyhexose)1 + (Man)3(GlcNAc)2 | 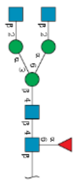 | 0.14% |

| | | | |
|---|---|---|---|
| G02763QD | (Hex)1 + (Man)3(GlcNAc)2 | 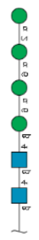 | 0.13% |
| G64481DJ | (HexNAc)1 + (Man)3(GlcNAc)2 | 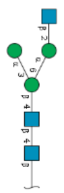 | 0.11% |
| G91365ZQ | (Hex)2 (HexNAc)2 (NeuAc)1 + (Man)3(GlcNAc)2 | 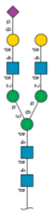 | 0.11% |
| G27919IH | (HexNAc)2 (Hex)1 (Deoxyhexose)1 + (Man)3(GlcNAc)2 | 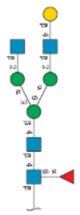 | 0.09% |
| G68668TB | (Hex)4 + (Man)3(GlcNAc)2 | 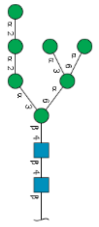 | 0.09% |
| G92024SL | (HexNAc)2 + (Man)3(GlcNAc)2 | 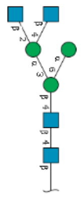 | 0.08% |

| | | | |
|---|---|---|---|
| G01760ZU | (Hex)1 + (Man)3(GlcNAc)2 | 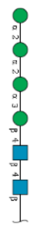 | 0.04% |
| G61751GZ | (HexNAc)1 + (Man)3(GlcNAc)2 | 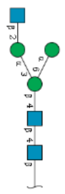 | 0.03% |
| G83161QT | (Hex)4 + (Man)3(GlcNAc)2 | 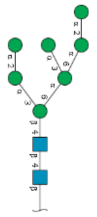 | 0.03% |
| G00176HZ | (HexNAc)4 (Deoxyhexose)1 + (Man)3(GlcNAc)2 | 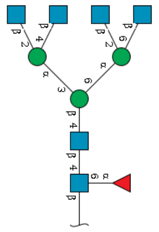 | 0.02% |
| G40358EZ | (HexNAc)5 (Deoxyhexose)1 + (Man)3(GlcNAc)2 | 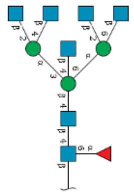 | 0.02% |
| G45560HM | (Hex)2 (HexNAc)2 (NeuAc)1 + (Man)3(GlcNAc)2 | 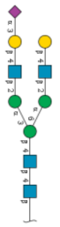 | 0.02% |
| G82348BZ | (Deoxyhexose)1 + (Man)3(GlcNAc)2 | 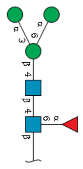 | 0.02% |

| | | | |
|---|---|---|---|
| G17473NR | (Hex)2 (HexNAc)3 (NeuAc)2 (Deoxyhexose)1 + (Man)3(GlcNAc)2 | 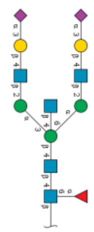 | 0.01% |
| G38117WE | (Hex)1 (HexNAc)2 (NeuAc)1 (Deoxyhexose)1 + (Man)3(GlcNAc)2 | 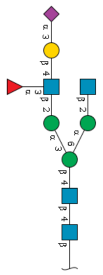 | 0.01% |
| G50078RT | (HexNAc)4 (Hex)3 + (Man)3(GlcNAc)2 | 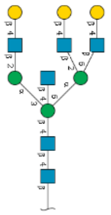 | 0.01% |
| G84467IZ | (Hex)2 (HexNAc)2 (NeuAc)2 + (Man)3(GlcNAc)2 | 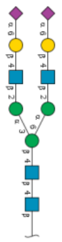 | 0.01% |
| G07483YN | (HexNAc)1 (Deoxyhexose)1 + (Man)3(GlcNAc)2 | 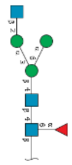 | 0.00% |
| G08997GC | (Hex)2 (HexNAc)2 (NeuAc)2 + (Man)3(GlcNAc)2 | 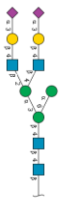 | 0.00% |

| | | | |
|---|---|---|---|
| G39213VZ | (HexNAc)2 + (Man)3(GlcNAc)2 | 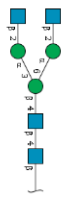 | 0.00% |

**Supplementary Figure 1: Number of GPSMs Found by Fraction.**

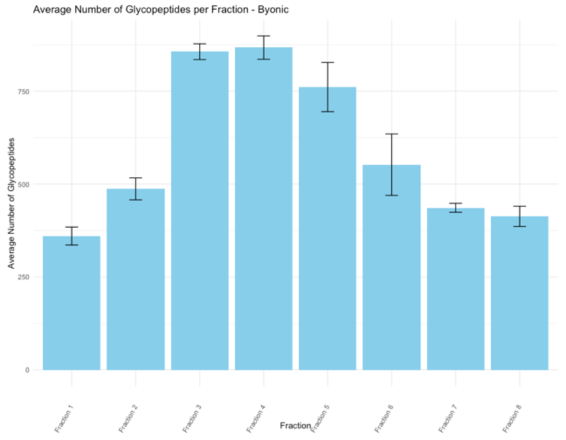

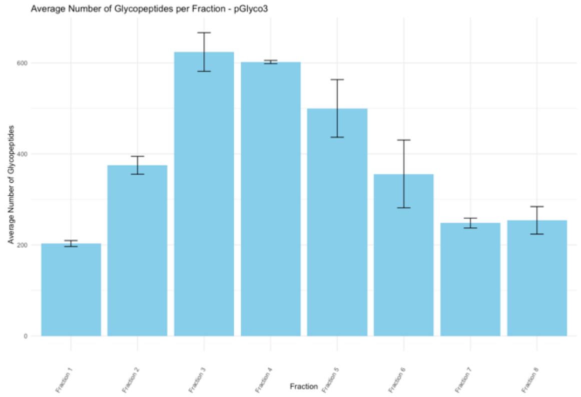
Average Number of Glycopeptides per Fraction - pGlyco3
Average Number of Glycopeptides
Fraction
Fraction 1
Fraction 2
Fraction 3
Fraction 4
Fraction 5
Fraction 6
Fraction 7
Fraction 8
0
200
400
600

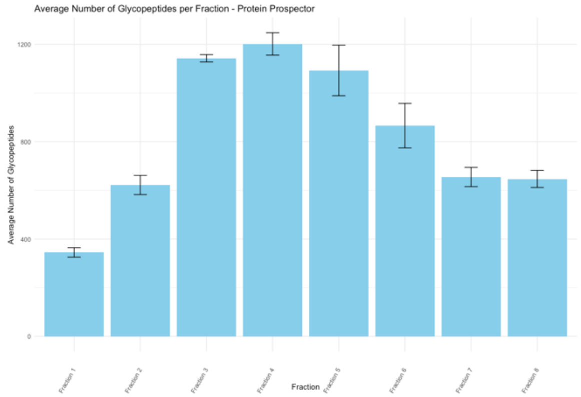
Average Number of Glycopeptides per Fraction - Protein Prospector
Average Number of Glycopeptides
Fraction
Fraction 1
Fraction 2
Fraction 3
Fraction 4
Fraction 5
Fraction 6
Fraction 7
Fraction 8
0
400
800
1200

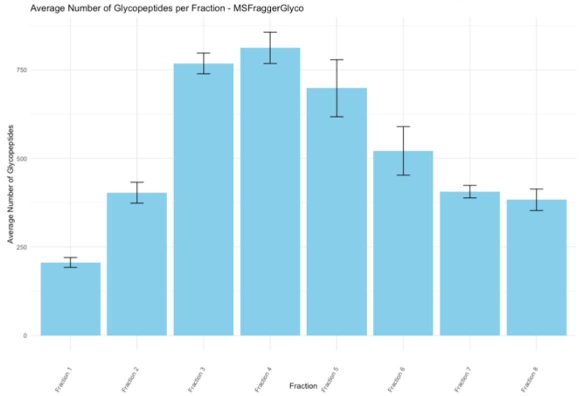
Average Number of Glycopeptides per Fraction - MSFraggerGlyco
Average Number of Glycopeptides
Fraction
Fraction 1
Fraction 2
Fraction 3
Fraction 4
Fraction 5
Fraction 6
Fraction 7
Fraction 8
0
250
500
750

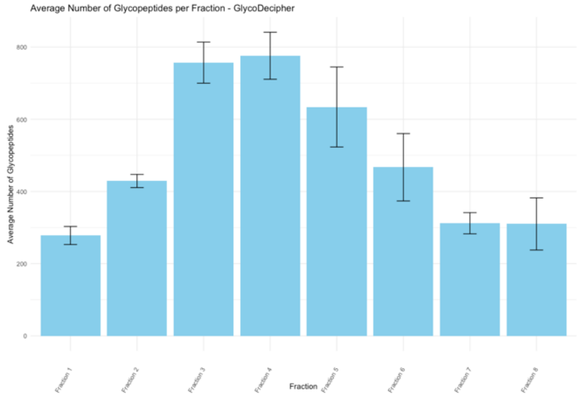
Average Number of Glycopeptides per Fraction - GlycoDecipher
Average Number of Glycopeptides
Fraction
Fraction 1
Fraction 2
Fraction 3
Fraction 4
Fraction 5
Fraction 6
Fraction 7
Fraction 8
0
200
400
600
800

**Supplementary Figure 2: Annotated spectra for five glycoforms of the peptide VQGNSTLLHITDLQAR from Immunoglobulin superfamily member 3.** Only the mass region from *m/z* 400 up is shown to spread out peaks and reduce overlaying of assignments (prominent HexNAc and HexNAcHex oxonium ions were seen in the missing low mass region for all these spectra). The $HexNAc_5Hex_4Fuc$ glycoform was identified by multiple software, but the other glycoforms were only reported by Protein Prospector.

**VQGN(HexNAc5Hex4Fuc*)STLLHITDLQAR$^{+3}$**

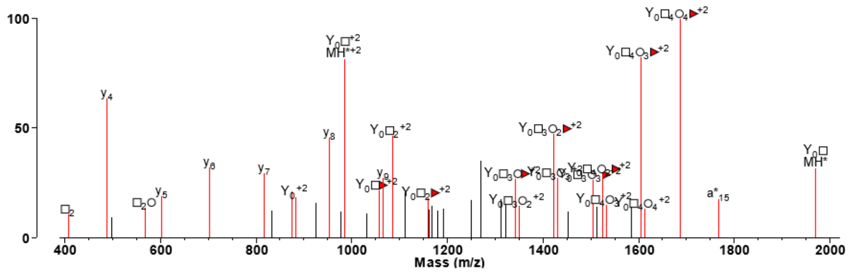


**VQGN(HexNAc4Hex4*)STLLHITDLQAR$^{+3}$**

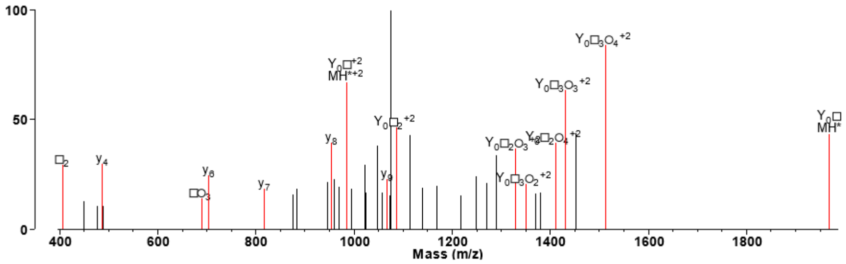


**VQGN(HexNAc4Hex4Fuc*)STLLHITDLQAR$^{+3}$**

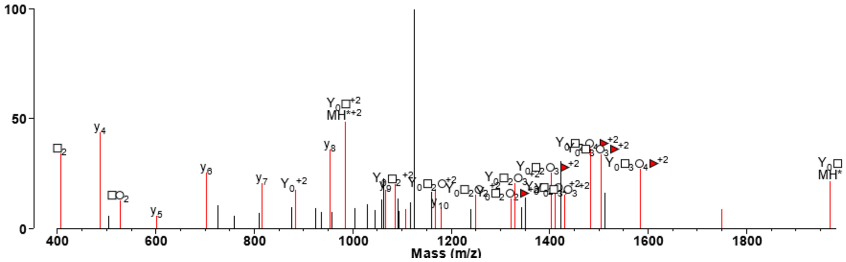


**VQGN(HexNAc5Hex4*)STLLHITDLQAR$^{+3}$**

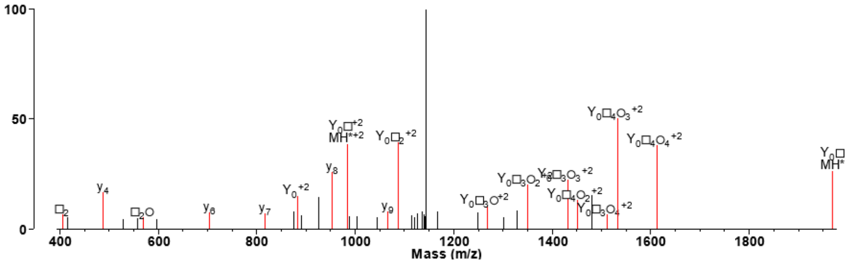


**VQGN(HexNAc2Hex5*)STLLHITDLQAR$^{+3}$**

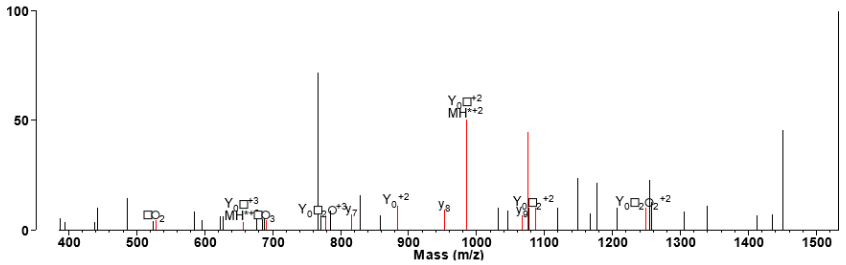

**Supplementary Figure 3: UniProtIDs with the Most Glycosites as Determined in Each Software.**

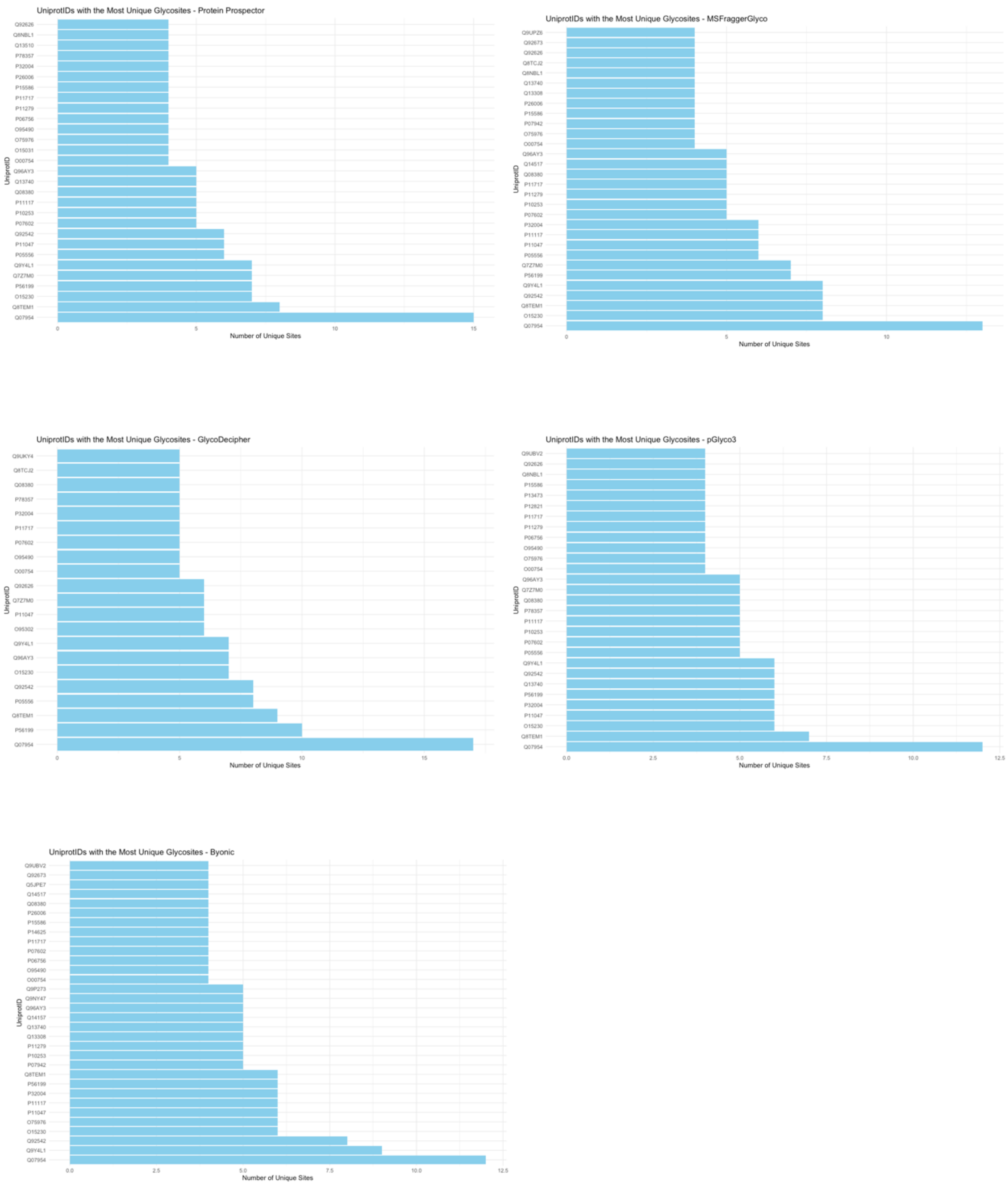

**Supplementary Figure 4: Percentage of Multi-Fucosylated Glycopeptides by Software**

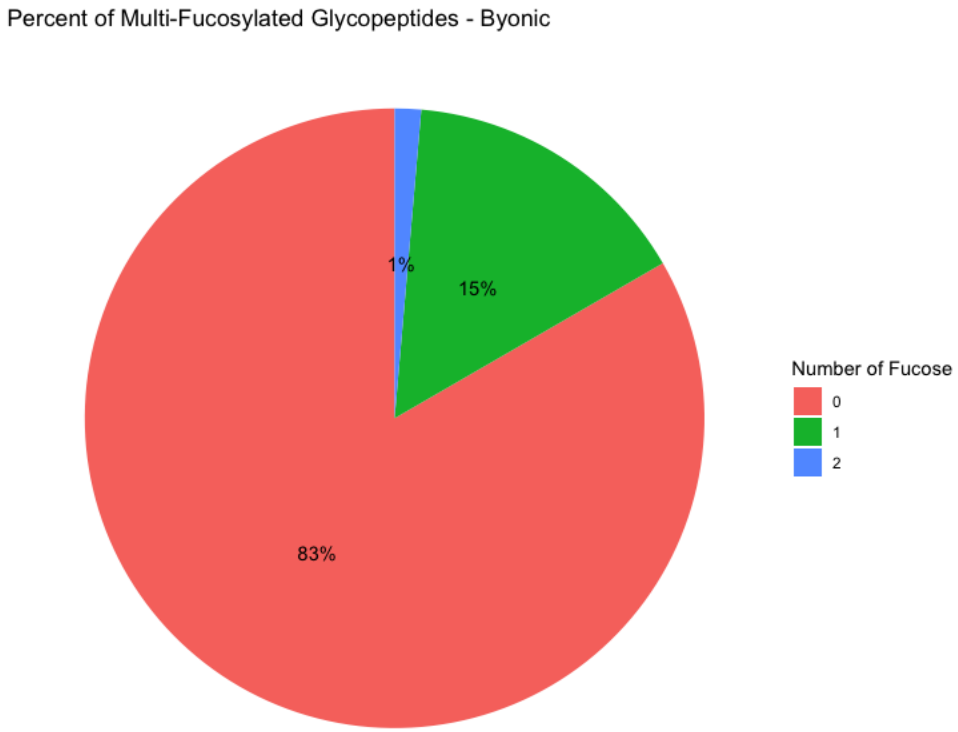


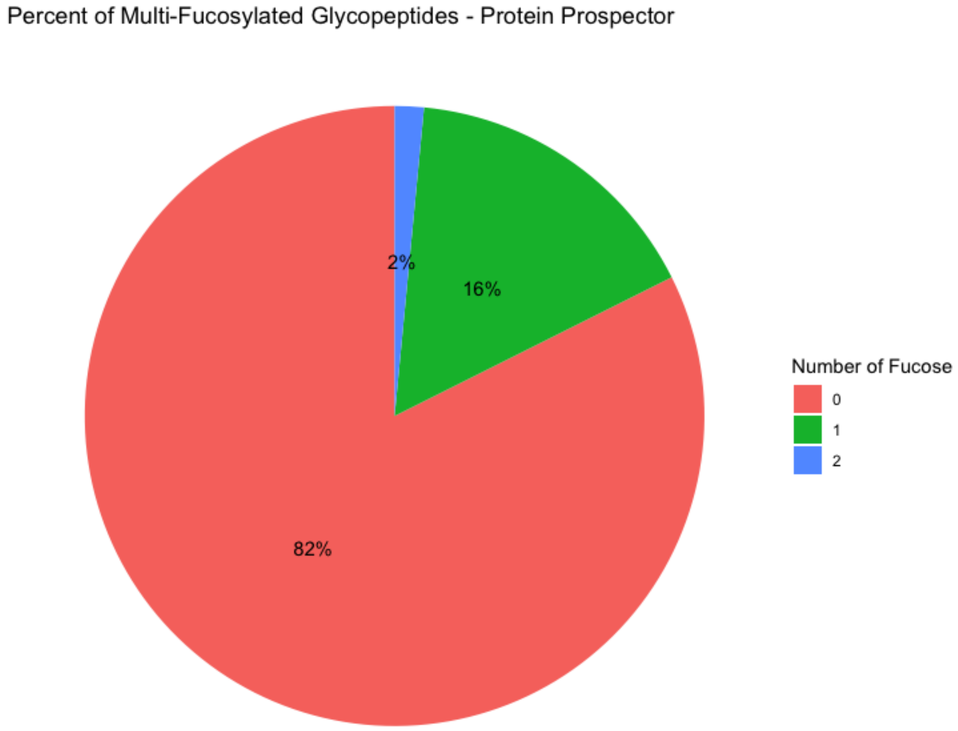

Percent of Multi-Fucosylated Glycopeptides - MSFraggerGlyco

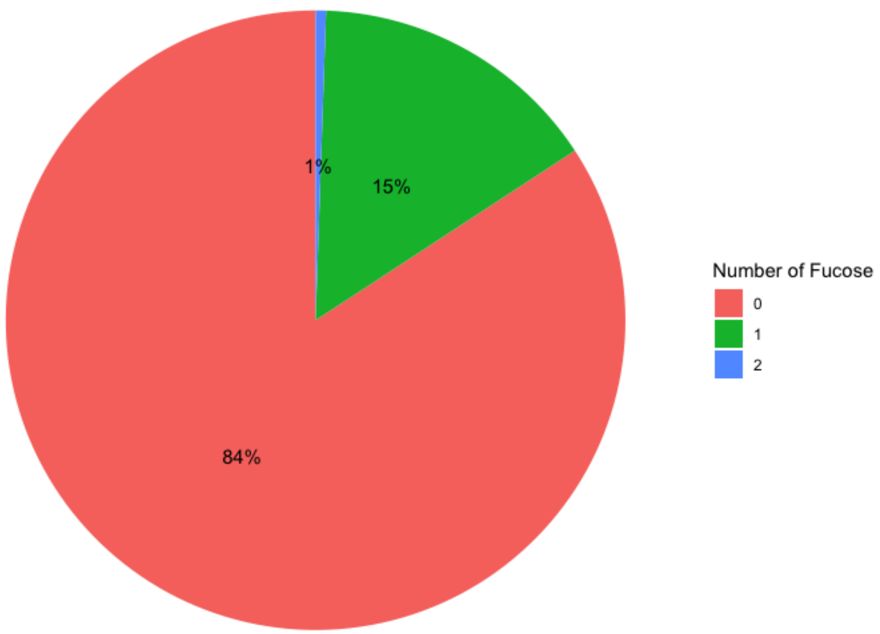


Percent of Multi-Fucosylated Glycopeptides - pGlyco3

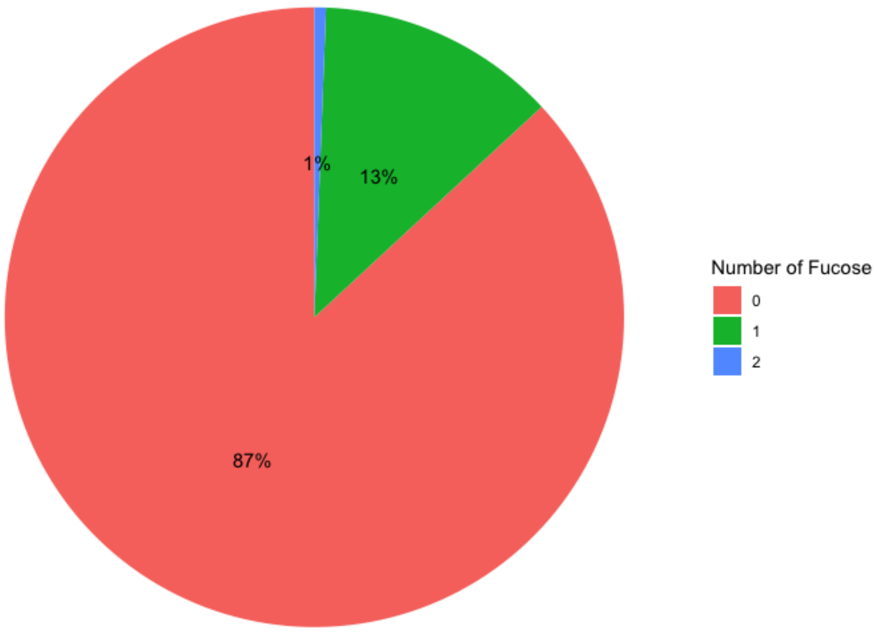

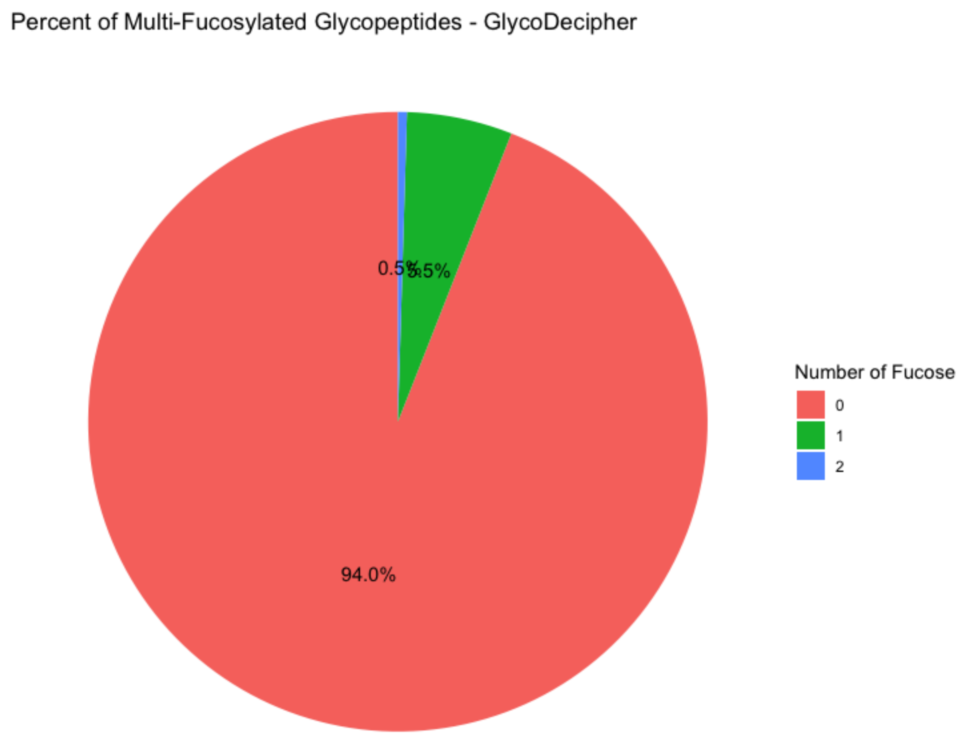
Percent of Multi-Fucosylated Glycopeptides - GlycoDecipher
0.5%
5.5%
94.0%
Number of Fucose
0
1
2

**Supplementary Figure 5**: **Speed of Each Glycoproteomic Search**

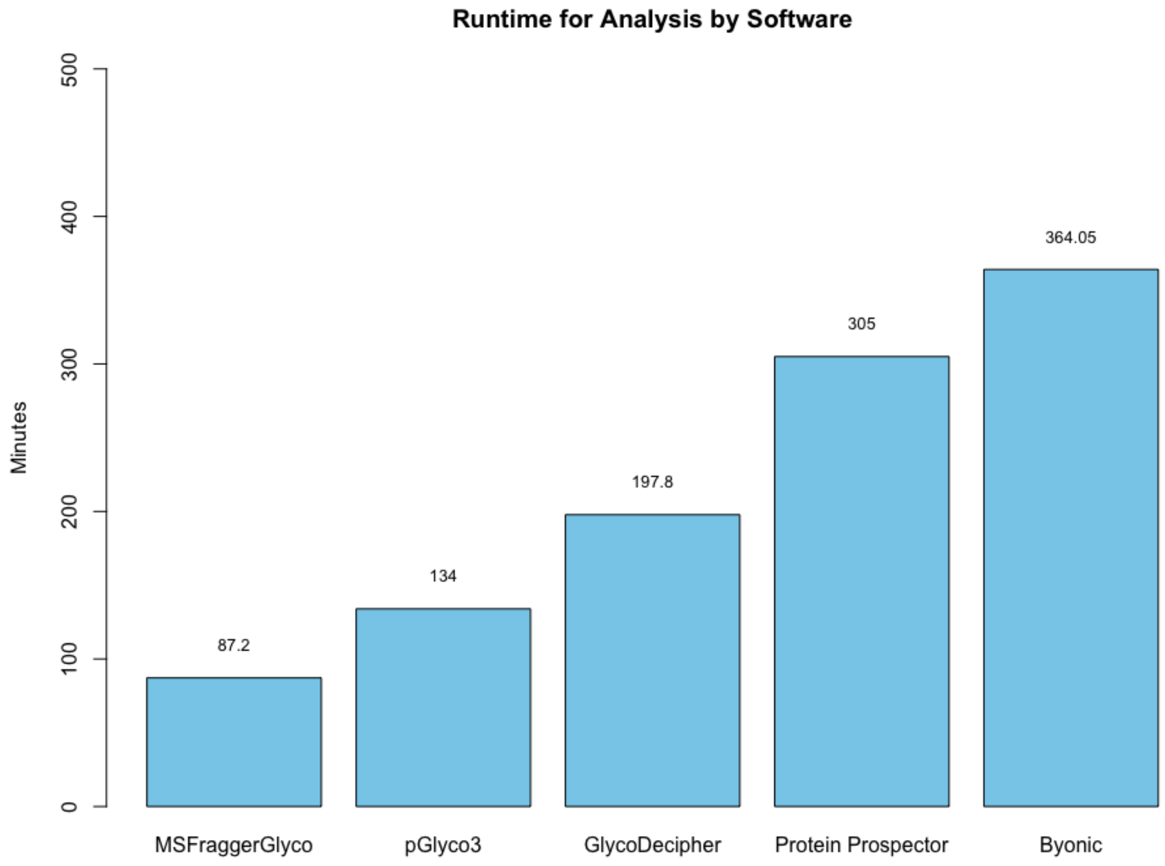

**Supplementary Figure 6**: **Mass Error of Peptides Detected in Byonic**

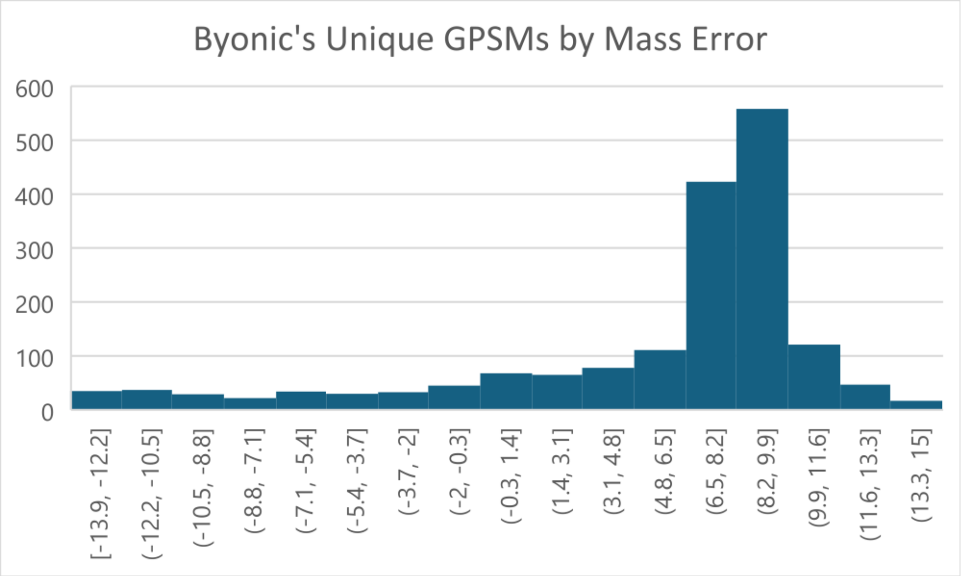


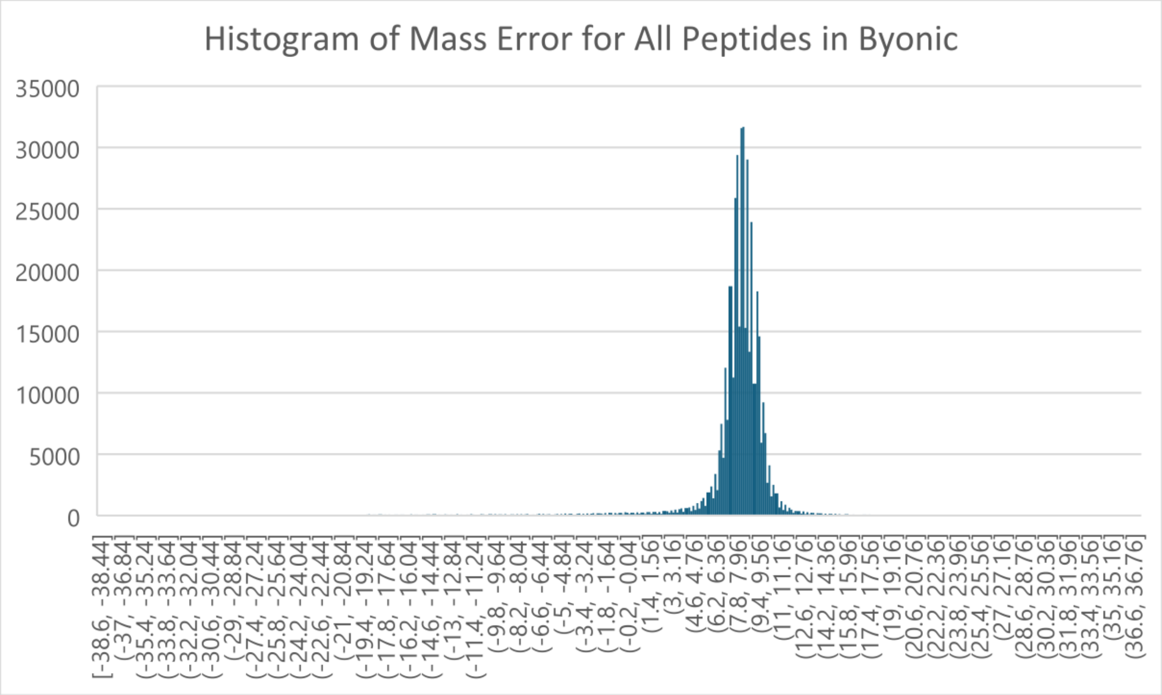
